# mRNA Location and Translation Rate Determine Protein Targeting to Dual Destinations

**DOI:** 10.1101/2023.04.24.538105

**Authors:** Alexander N. Gasparski, Konstadinos Moissoglu, Sandeep Pallikkuth, Sezen Meydan, Nicholas R. Guydosh, Stavroula Mili

**Author notes:** Correspondence, Stavroula Mili, Laboratory of Cellular and Molecular Biology NCI, NIH, 37 Convent Drive, MSC 4256, Building 37, Room 2042, Bethesda, MD 20892-4256.

## Abstract

Numerous proteins are targeted to two or multiple subcellular destinations where they exert distinct functional consequences. The balance between such differential targeting is thought to be determined post-translationally, relying on protein sorting mechanisms. Here, we show that protein targeting can additionally be determined by mRNA location and translation rate, through modulating protein binding to specific interacting partners. Peripheral localization of the *NET1* mRNA and fast translation lead to higher cytosolic retention of the NET1 protein, through promoting its binding to the membrane-associated scaffold protein CASK. By contrast, perinuclear mRNA location and/or slower translation rate favor nuclear targeting, through promoting binding to importins. This mRNA location-dependent mechanism is modulated by physiological stimuli and profoundly impacts NET1 function in cell motility. These results reveal that the location of protein synthesis and the rate of translation elongation act in coordination as a ‘partner-selection’ mechanism that robustly influences protein distribution and function.

## Introduction

Numerous and diverse eukaryotic proteins are targeted to two or multiple subcellular destinations (Karniely and Pines, 2005). Proteins with dual localization include metabolic enzymes, signaling factors and apoptosis regulators (Gough et al., 2009; Mihara et al., 2003; Popgeorgiev et al., 2018; Wegrzyn et al., 2009; Yogev et al., 2011). Alterations in the balance of protein targeting affect important physiological outcomes (De Jesus et al., 2022; Knockaert et al., 2011; Raza, 2011). Dual targeting is often regulated through post-translational mechanisms that favor the action of one out of two or more competing targeting signals or promote processes or interactions that retain the polypeptide in a specific compartment (Avadhani, 2011; Avadhani et al., 2011; Karniely and Pines, 2005).

NET1 is a guanine nucleotide exchange factor, which activates the RhoA GTPase, and is distributed in both the nucleus and cytoplasm. Cytoplasmic NET1 regulates RhoA and controls the cytoskeleton and cell migration, while nuclear NET1 is thought to control cellular responses to DNA damage (Dubash et al., 2011; Schmidt and Hall, 2002; Srougi and Burridge, 2011). Post-translational modifications can alter NET1 distribution and function (Song et al., 2015; Ulu et al., 2018). Interestingly however, NET1 gene expression exhibits an additional level of regulation at the level of mRNA localization. The *NET1* mRNA is prominently targeted to peripheral protrusive cytoplasmic regions of migrating cells (Mili et al., 2008; Wang et al., 2017). While compartmentalized distribution in the cytosol has now been described for a large fraction of mammalian mRNAs, the precise functional consequences of this prevalent mode of regulation are poorly understood (Das et al., 2021; Gasparski et al., 2022). Peripheral *NET1* mRNA localization is observed in *in vivo* tumors and is important for cancer cell invasion (Chrisafis et al., 2020). However how peripheral mRNA localization and translation influence NET1 protein function is unknown. Here, we show that *NET1* mRNA location in combination with the rate of its translation can influence the ability of competing domains to determine the targeting of the newly synthesized protein, through favoring binding to specific partners. This RNA-based ‘partner-selection’ mechanism provides a robust means of controlling protein function.

## Results

### Altering NET1 mRNA location between peripheral and perinuclear regions

Localization of the *NET1* mRNA to the cell periphery is an active process that requires sequences present in the 3’UTR of the transcript. Specifically, NET1 and co-regulated mRNAs are trafficked to the periphery through the KIF1C kinesin, and contain GA-rich regions in their 3’UTRs, which are necessary for localization (Chrisafis *et al*., 2020; Moissoglu et al., 2020; Pichon et al., 2021). Antisense phosphorodiamidate morpholino oligonucleotides (PMOs) specific to these regions can interfere with mRNA targeting (Chrisafis *et al*., 2020; Moissoglu *et al*., 2020). We tiled 25nt-long PMOs across most of the length of the human *NET1* 3’UTR (Figure S1A) and tested for their ability to interfere with peripheral *NET1* mRNA localization (Figure S1B) as well as for their effects on overall *NET1* mRNA and NET1 protein levels (Figure S1C-D). This allowed us to identify PMOs that result in an almost diffuse *NET1* mRNA distribution (Figure S1B; indicated by a PDI (Peripheral Distribution Index) of 1; (Stueland et al., 2019)) and which have no apparent effect on *NET1* mRNA levels or translation efficiency. Consistent with prior observations (Chrisafis *et al*., 2020), targeting the region highest in GA-content with a combination of two PMOs (921 and 975) significantly reduced peripheral *NET1* mRNA localization (Figures 1A, S1B, S2A). Additional PMOs in distant regions produced a similar effect, suggesting the presence of multiple necessary elements or the involvement of long-range, tertiary interactions. Regardless of the exact mechanism, PMOs^921+975^ and PMO^1067^ provided us with tools that allow the conversion of the *NET1* mRNA distribution from a peripheral one to mostly diffuse or perinuclear (Figure 1A and S2A). Significantly, this alteration was not accompanied by changes in *NET1* mRNA or protein levels (Figure 1B and S2B) and was specific for NET1 since the distribution of the co-regulated *RAB13* mRNA was not impacted (Figure S2C).

**Figure 1:**
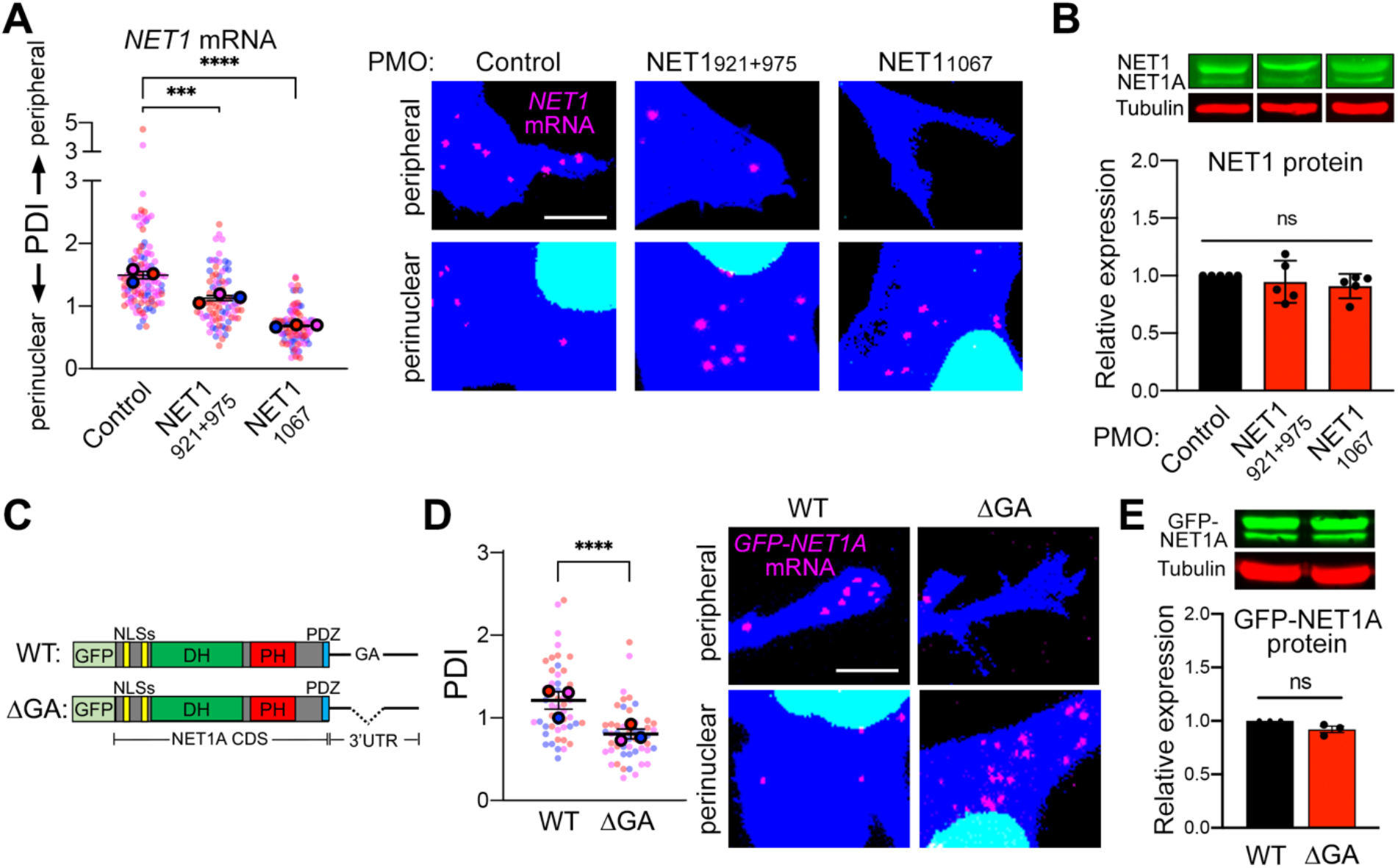
Altering *NET1* mRNA localization between peripheral and perinuclear regions. **(A)** FISH images of MDA-MB-231 cells treated with the indicated PMOs. Zoomed in perinuclear and peripheral regions are shown (whole images are shown in Figure S2A). The graph presents PDI quantification of *NET1* mRNA distribution, with higher values indicating a more peripheral mRNA distribution. PDI=1 indicates a random, diffuse distribution. n=84-118 cells from 3 independent experiments. **(B)** Protein levels of NET1 and NET1A isoforms, by Western blot, in cells treated with the indicated PMOs. n=5. **(C)** Schematic of the GFP-NET1A constructs used for generation of stably expressing cell lines. Coding sequences (CDS) are identical. The ΔGA construct carries a 78nt deletion of a GA-rich region within the 3’UTR. **(D)** FISH images and corresponding PDI values of GFP-NET1A mRNA in WT and ΔGA expressing cell lines. Zoomed in perinuclear and peripheral regions are shown (whole images are shown in Figure S2D). n=50 cells from 3 independent experiments. **(E)** GFP-NET1A protein levels, by Western blot, in WT and ΔGA expressing cell lines. n=3. In superplots, data points from individual replicates are color coded, and large, outlined color dots indicate the mean of each replicate. Error bars: SEM. p-values: ***<0.001, ****<0.0001, ns: not significant by one-way ANOVA (A, B) or unpaired t-test (D, E). Scale bars: 4μm.

To independently manipulate *NET1* mRNA localization, we generated stable cell lines that express constructs encoding a fusion protein of GFP with NET1A, the NET1 isoform mostly implicated with cell motility phenotypes (Carr et al., 2013; Song *et al*., 2015). The sequence encoding this fusion protein was attached to either the full-length wild type (WT) *NET1* 3’UTR, or a deletion mutant with the region highest in GA-content removed (ΔGA) (Figure 1C). Consistent with the above results, the ΔGA mutant mRNA exhibited a perinuclear distribution that was significantly different from the peripheral localization imparted by the WT 3’ UTR (Figure 1D and S2D). Importantly, the overall *GFP-NET1A* mRNA and GFP-NET1A protein levels produced from these constructs were indistinguishable (Figure 1E and S2E, F), suggesting that the *NET1* mRNA is equally stable and translated with similar efficiency in peripheral and perinuclear locations. Overall, PMO delivery or 3’UTR deletion specifically alters the distribution of endogenous or exogenous *NET1* mRNA respectively, and thus allows us to further understand the molecular and functional consequences imparted by changing the site of NET1 protein synthesis.

### mRNA location specifies NET1 protein targeting

The NET1 protein is dually targeted to both the nucleus and the cytosol. We thus tested whether altering *NET1* mRNA location impacts the overall targeting of the protein between these two destinations. We assessed, in live cells, the nucleo-cytoplasmic distribution of GFP-NET1A produced from either a peripheral mRNA (GFP-NET1A/WT UTR) or from a perinuclear mRNA, upon treatment with mis-localizing antisense PMOs (PMOs^921+975^ and PMO^1067^; Figure 2A) or deletion of the GA-rich region within the 3’ UTR (GFP-NET1A/ΔGA UTR; Figure 2B). Interestingly, perinuclearly produced NET1A partitioned to a higher extent within the nucleus (Figure 2A, B). Of note, the amount of total protein produced under all conditions is the same, as shown above (Figure 1B, E and S2F), indicating that the reduction in cytoplasmic protein is due to increased import into the nucleus and not due to increased degradation in the cytosol. Furthermore, nuclear accumulation is abolished when the two N-terminal basic NLSs are mutated (Figure 2B), suggesting that NET1A nuclear import involves the action of nuclear transport receptors of the importin/karyopherin superfamily.

**Figure 2:**
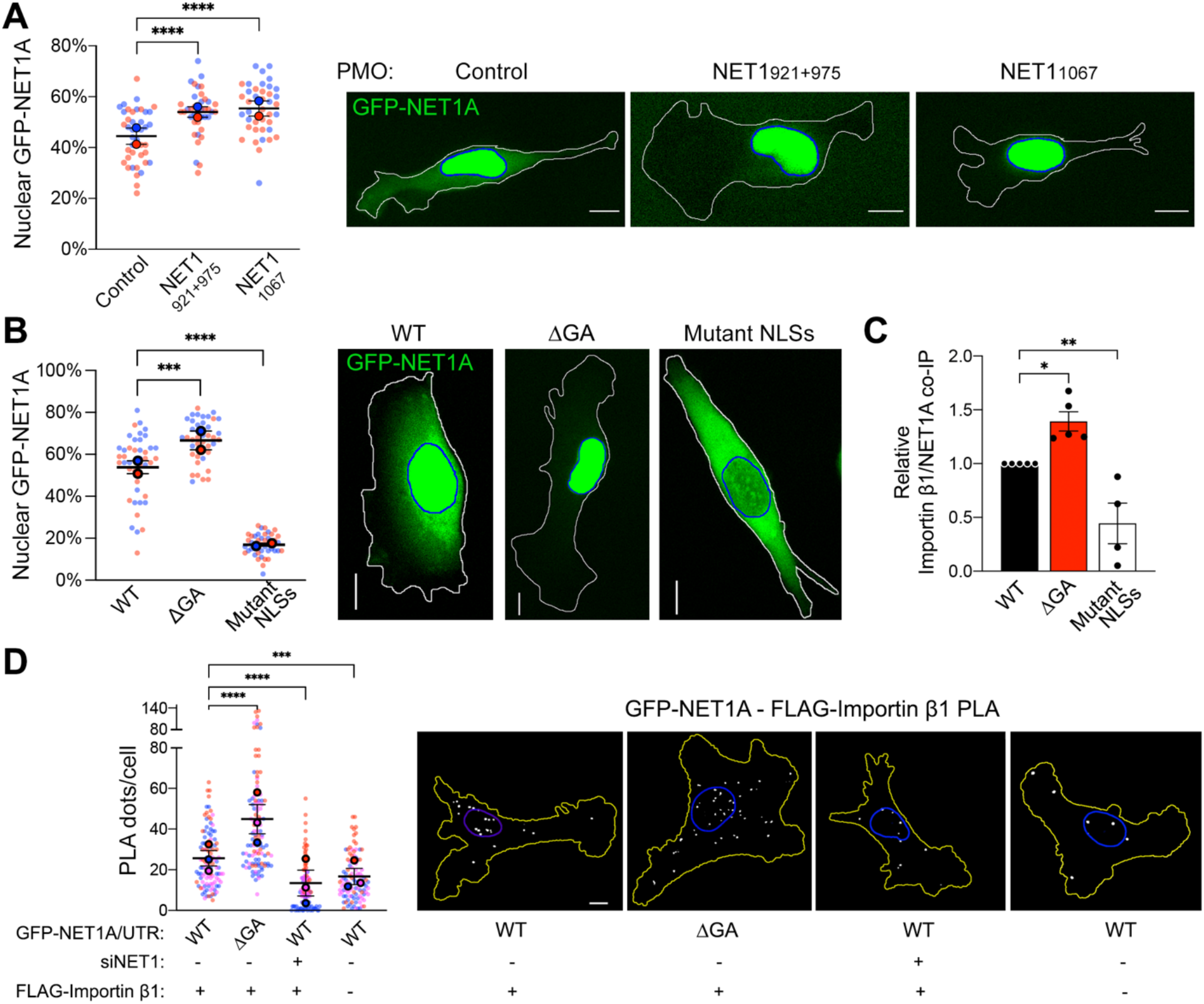
*NET1* mRNA location determines NET1-importin binding and nucleocytoplasmic distribution. **(A)** Live GFP fluorescence imaging of cells expressing GFP-NET1A/WT UTR treated with the indicated PMOs. The percent GFP-NET1A signal within the nucleus is quantified. n=39-41 in 2 independent experiments. **(B)** Live GFP fluorescence imaging of the indicated cell lines (see Figure 1C) and quantification of the percent GFP-NET1A signal within the nucleus. Blue outline: nuclear boundary; white outline: cell boundary. n=43-44 in 2 independent experiments. **(C)** Relative importin β1 binding to GFP-NET1A from co-immunoprecipitation (co-IP) experiments of the indicated cell lines. GFP trap beads were used for IP and GFP-NET1A and FLAG-importin β1 proteins were detected by Western blot. n=5. **(D)** In situ detection of interaction between GFP-NET1A and FLAG-importin β1, by PLA in the indicated cell lines. White dots: PLA signal; blue outline: nuclear boundary; yellow outline: cell boundary. n=104-113 in 3 independent experiments. In superplots, data points from individual replicates are color coded, and large, outlined color dots indicate the mean of each replicate. Error bars: SEM. p-values: *<0.05, **<0.01, ***<0.001, ****<0.0001, ns: not significant by one-way ANOVA. Scale bars: 10μm.

To test whether the site of NET1A synthesis affects its interaction with transport receptors, we assessed NET1A association with importin β1/KPNB1, the main transport receptor involved in import of basic NLSs, in a heterodimer with an importin α family member. We detected NET1A-importin β1 interaction by co-immunoprecipitation (Figure 2C) or proximity ligation amplification (PLA), which allows the detection of complex formation between two partners in situ (Figure 2D). Indeed, importin β1 specifically co-IPs with GFP-NET1A in a manner dependent on the presence of functional NLSs (Figure 2C). Furthermore, PLA can specifically report the in situ presence of NET1A-importin β1 complex, in both the cytosol and nucleus, since the observed signal is significantly reduced when either partner is knocked-down (Figure 2D; +siNET1) or missing (Figure 2D; -FLAG-importin β1). Both assays revealed that GFP-NET1A produced perinuclearly (ΔGA UTR) exhibits an increased interaction with importin β1, consistent with its higher nuclear partitioning. Of note, most of the NET1A-importin β1 complex is observed perinuclearly, in agreement with reports showing that importin α and β subunits are preferentially enriched towards the perinuclear cell body rather than peripheral protrusive regions (Mardakheh et al., 2015).

### Newly synthesized NET1 interacts with importin

The observation that the location of the *NET1* mRNA affected the degree with which the encoded NET1 protein interacted with importin β1, prompted us to address whether mRNA translation is also involved. We thus assessed NET1A-importin β1 interaction after a brief (20 min) treatment with cycloheximide or puromycin to block translation. Interestingly, translation inhibition abolished the increased interaction with importin β1 observed for perinuclearly produced NET1A (ΔGA UTR) (Figure 3A, B), suggesting that importin β1 largely interacts with newly synthesized NET1A. We interpret these data to indicate that perinuclearly synthesized NET1A rapidly and preferentially interacts with nuclear import receptors. Given the presence of NLSs at the very N-terminus this interaction could likely occur co-translationally. Import into the nucleus would then result in dissociation of the transport receptors. By contrast, peripherally synthesized NET1A interacts with importin β1 less efficiently and independent of active translation, potentially reflecting a low level of nucleocytoplasmic shuttling of mature NET1A protein.

**Figure 3:**
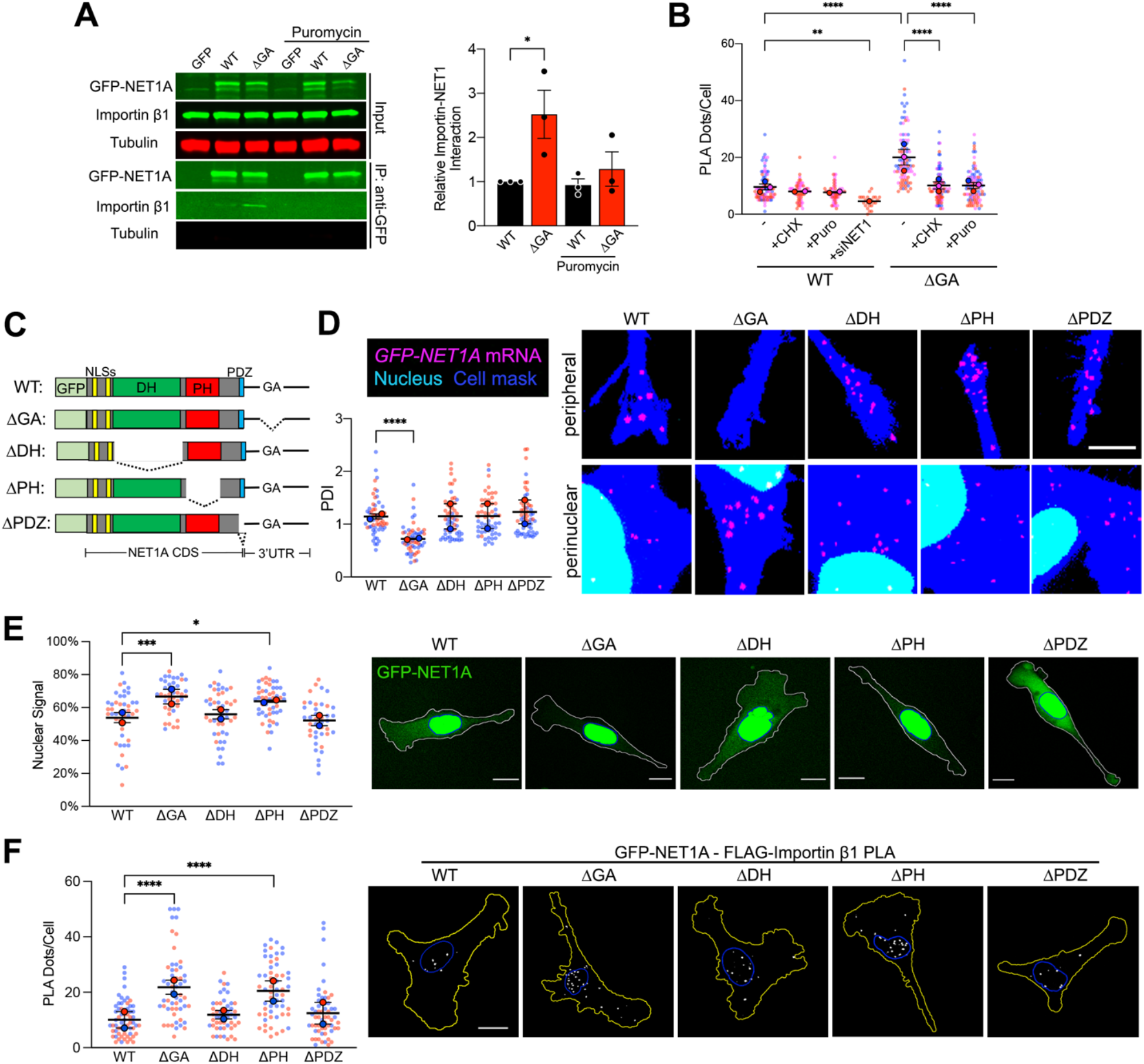
The NLS and PH domains competitively determine nuclear import of newly-synthesized NET1A protein. **(A)** Importin β1 binding to GFP or GFP-NET1A from co-immunoprecipitation experiments of the indicated MDA-MB-231 cell lines with 20min puromycin treatment. Quantifications are shown relative to the WT untreated sample. n=3. **(B)** Quantification of in situ interaction between GFP-NET1A and FLAG-importin β1, by PLA of the indicated cell lines, with 20min cycloheximide (CHX) or puromycin (Puro) treatment, or NET1 knockdown (siNET1). n=55-98 in 2-3 independent experiments. For siNET1 n=25. **(C)** Schematic of the GFP-NET1A constructs used for generation of stably expressing MDA-MB-231 cell lines. Dotted lines indicate deleted regions. **(D)** FISH images and corresponding PDI values of GFP-NET1A mRNA distribution in the indicated stable cell lines. Zoomed in perinuclear and peripheral regions are shown. n=55-58 cells from 2 independent experiments. **(E)** Live GFP fluorescence imaging of the indicated cell lines and quantification of the percent GFP-NET1A signal within the nucleus. Blue outline: nuclear boundary; white outline: cell boundary. n=40-50 in 2 independent experiments. **(F)** In situ detection of interaction between GFP-NET1A and FLAG-importin β1, by PLA in the indicated cell lines. White dots: PLA signal; blue outline: nuclear boundary; yellow outline: cell boundary. n=49-59 in 2 independent experiments. In superplots, data points from individual replicates are color coded, and large, outlined color dots indicate the mean of each replicate. Error bars: SEM. p-values: *<0.05, **<0.01, ***<0.001, ****<0.0001 by unpaired t-test (A) or one-way ANOVA (B, D, E, F). Scale bars: 4μm (D); 15μm (E, F).

### Competition between NET1 protein domains specifies targeting

Polypeptides that can be targeted to dual destinations often contain distinct signals or domains that direct targeting to one or the other location. In the case of NET1A, as detailed above, nuclear targeting is mediated through the action of the N-terminal NLSs. Other recognizable domains could function to promote cytoplasmic retention. These include: a Dbl-homology (DH) domain, characteristic of proteins that activate Rho family GTPases by functioning as guanine nucleotide exchange factors (GEFs); a Plekstrin-homology (PH) domain, usually involved in protein-protein or protein-membrane interactions; and a C-terminal PDZ motif recognized by PDZ domain-containing partners (Figure 3C). To determine the involvement of these domains in cytosolic retention of NET1A, we generated stable cell lines expressing deletion mutants (Figure 3C and S3). All mutants were expressed from constructs carrying the WT NET1 UTR, which as expected resulted in peripheral mRNA localization similar to that observed for the full-length protein (Figure 3D). Deletion of the DH domain or of the PDZ motif did not alter NET1A association with importin β1, or the fraction partitioning in the nucleus (Figure 3E, F). Interestingly, however, deletion of the PH domain promoted NET1A-importin β1 interaction and increased nuclear accumulation to a degree similar to that seen when full-length NET1A is produced from a perinuclear mRNA (Figure 3E, F). Therefore, the PH domain appears to be in competition with the NLSs to determine the eventual nucleo-cytoplasmic balance of NET1A targeting. Given that deletion of the DH domain does not exhibit a similar effect, we conclude that the GEF activity *per se* is not necessary for cytoplasmic retention, but that rather some other factor recognized by the PH domain functions to suppress NET1A nuclear import.

### NET1 mRNA location determines protein targeting through partner selection

To identify this factor, we sought, through mass spectrometry analysis, for GFP-NET1A binding partners whose interaction is promoted by peripheral localization of the *NET1* mRNA. We focused on one candidate, the membrane-associated scaffold protein CASK (LaConte et al., 2016; Porter et al., 2019) (Figure S4). Indeed, CASK interacts with GFP-NET1A in co-immunoprecipitation experiments and this interaction is minimized when NET1A is produced perinuclearly (Figure 4A; ΔGA). Furthermore, CASK-NET1A interaction requires the PH domain of NET1A (Figure 4A; ΔPH). The CASK-NET1A interaction can also be visualized by PLA which further revealed that the complex resides to a large extent close to the cell periphery (potentially reflecting membrane association) and verified that its formation depends on the location of the *NET1* mRNA (Figure 4B). Overall, the NET1A-CASK complex is regulated by the location of the *NET1* mRNA in a manner opposite to the NET1A-importin β1 complex and resides in spatially distinct regions in the cell.

**Figure 4:**
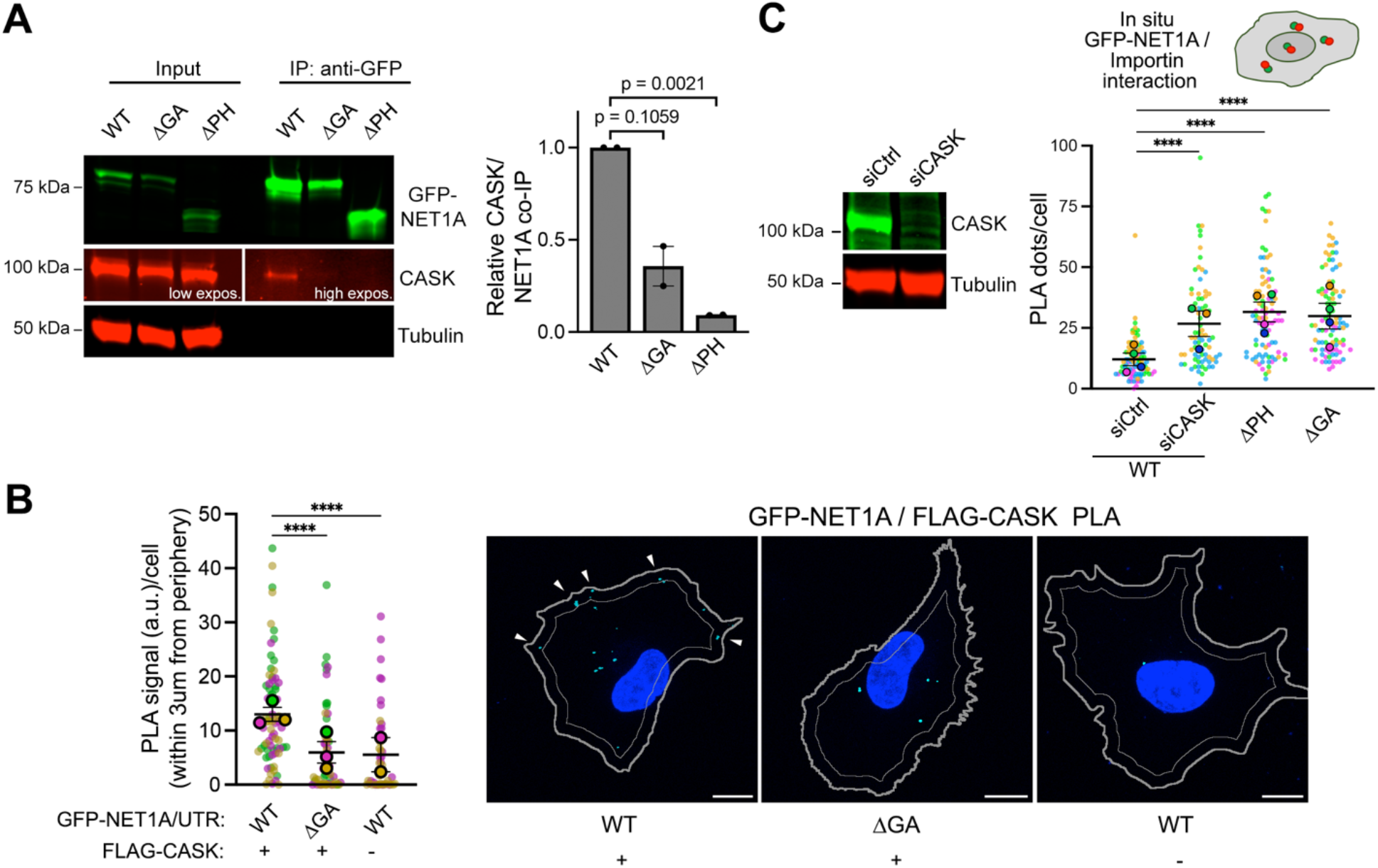
NET1A-CASK interaction competes with NET1A-importin binding and is regulated in an opposite manner by *NET1* mRNA location. **(A)** Representative Western blot and quantification of relative CASK binding to GFP-NET1A from co-immunoprecipitation experiments of the indicated MDA-MB-231 cell lines. n=2. **(B)** In situ detection of interaction between GFP-NET1A and FLAG-CASK, by PLA in the indicated cell lines. Cyan dots: PLA signal; blue: DAPI; thick gray outline: cell boundary; thin gray outline: inner boundary 3μm from periphery. n=45-72 in 3 independent experiments. **(C)** Western blot showing efficiency of CASK knockdown upon siRNA treatment and quantification of in situ interaction between GFP-NET1A and FLAG-importin β1, by PLA of the indicated cell lines (see figure 3C). n=78-92 in 4 independent experiments. In superplots, data points from individual replicates are color coded, and large, outlined color dots indicate the mean of each replicate. Error bars: SEM. p-values:, ****<0.0001 by one-way ANOVA (A, C) or Kruskal-Wallis test (B). Scale bars: 10μm.

To directly test whether CASK is the factor that suppresses importin β1 binding and NET1A nuclear import, we knocked-down CASK expression with siRNAs. Indeed, absence of CASK led to increased NET1A-importin β1 interaction, even though the *NET1* mRNA is peripheral, and mimicked the effect seen upon deletion of the PH domain or upon expression of NET1A from perinuclear mRNA (Figure 4C).

### Modulating the translation elongation rate through the NET1A coding sequence

Since recognition of the NLSs by importins occurs during, or shortly after, translation (Figure 3A, B), we reasoned that the ability of CASK to antagonize with NLS-importin binding would likely occur in a similar time frame. In this regard, the rate of translation elongation could influence the kinetics of appearance and/or folding of competing domains and thereby modulate their ability to direct eventual protein targeting. To assess the potential existence of any distinctive features associated with *NET1* mRNA translation, we examined monosome and disome ribosome footprint profiling data from human HEK293 cells (Meydan and Guydosh, 2020). These cells also exhibit peripheral *NET1* mRNA localization that is dependent on GA-rich sequences in the 3’UTR (Figure S5). Intriguingly, we observed higher monosome peaks, believed to reflect slower local elongation rates (Brar and Weissman, 2015), at two sites within the region encoding the DH domain (Figure 5A, arrows). These monosome peaks are also observed in other conditions and backgrounds (Figure S6A) and are detected around proline codons (Figure 5A), which have been described as a major contributor to ribosome stalling (Choi et al., 2018). Disome profiling revealed disome peaks upstream of these high monosome density sites (Figure 5A, S6B), further supporting the notion that they reflect regions of slower elongation leading to ribosome pile-up at upstream sequences.

**Figure 5.**
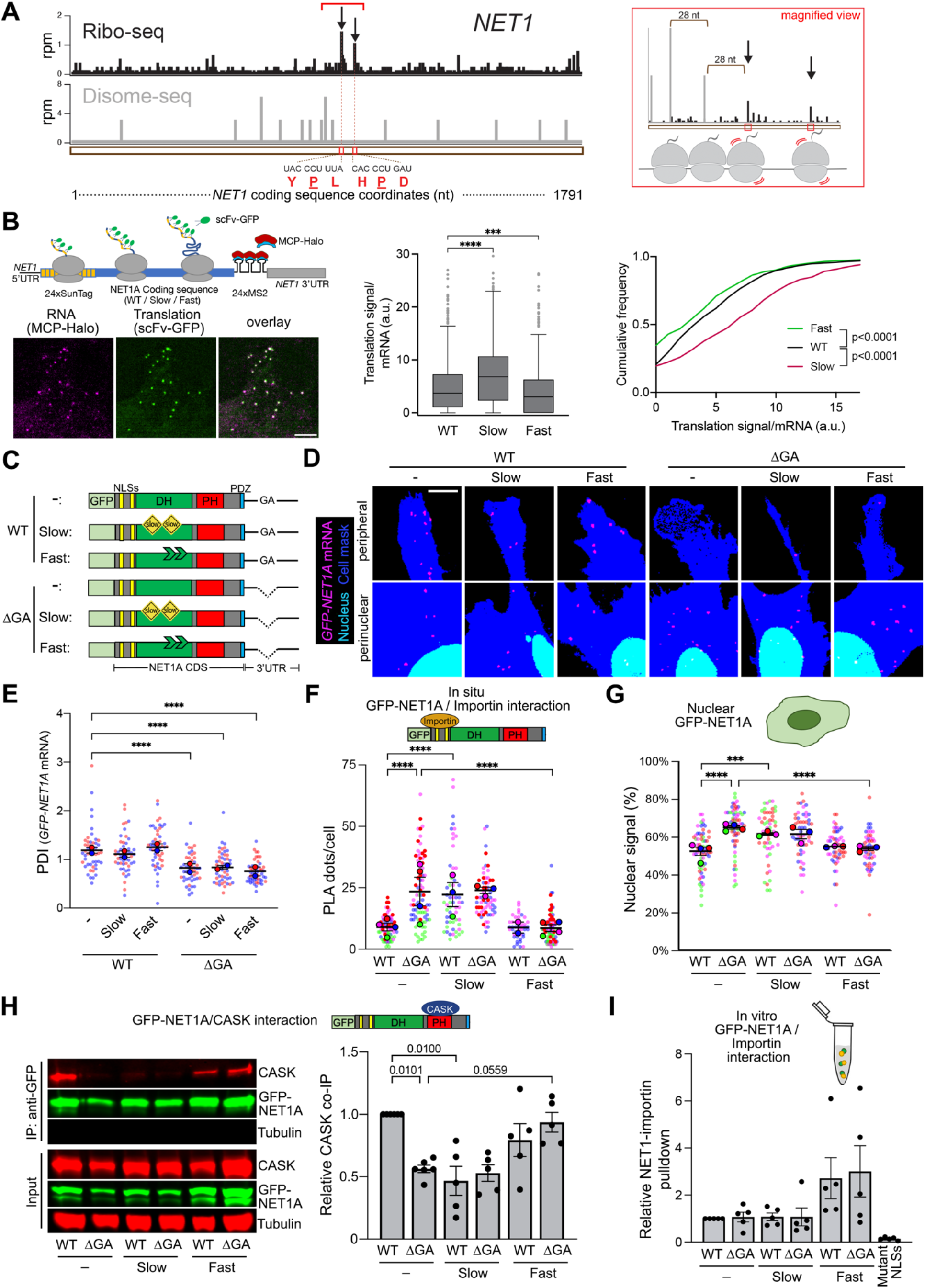
The rate of translation elongation determines partner selection and NET1 protein targeting. **(A)** Left panel: Snapshot of Ribo-seq (black) and Disome-seq (gray) reads mapped to the *NET1* transcript. For Ribo-seq, reads are plotted to correspond to the approximate P-site of the ribosome. For Disome-seq, reads are plotted to correspond to the approximate P-site of the trailing ribosome in the disome complex. Stalling peaks are indicated with arrows. The amino acid and nucleotide sequence of the stalling motifs corresponding to E, P, A sites of the ribosome are shown below the snapshot, with the P site amino acids underlined. Right panel: magnified snapshot of the Ribo-seq (black) and Disome-seq (gray) peaks within the region indicated by a red bracket in the left panel. Disome peaks ∼28nt (a typical ribosome footprint length) apart from each other and ∼28nt upstream of the first stalling site indicate the possibility of queued ribosomes in this region. **(B)** Schematic of NET1-based reporter constructs for single molecule translation site imaging. Examples of typical imaging snapshots are shown (left panel). Graphs present Tukey’s box-plot distributions (middle panel) or cumulative frequency plots (right panel) of translation signals/mRNA of the indicated constructs. n=604-863 mRNA particles in 40-42 cells. **(C)** Schematic of the GFP-NET1A constructs used for generation of stably expressing MDA-MB-231 cell lines. ‘Slow’ constructs include multiple suboptimal codons within the DH domain. ‘Fast’ constructs include mutations of the identified stall sites, also within the DH domain. **(D, E)** FISH images (D) and corresponding PDI values (E) of GFP-NET1A mRNA distribution in the indicated stable cell lines. Zoomed in perinuclear and peripheral regions are shown. n=46-52 cells from 2 independent experiments. **(F)** Quantification of in situ interaction between GFP-NET1A and FLAG-importin β1, by PLA of the indicated cell lines. n=44-86 in 3-4 independent experiments. **(G)** Quantification of the percent GFP-NET1A signal within the nucleus from live GFP fluorescence imaging of the indicated cell lines. n=51-65 in 3-4 independent experiments. **(H)** Representative Western blot and quantification of relative CASK binding to GFP-NET1A from co-immunoprecipitation experiments of the indicated cell lines. n=5. **(I)** Quantification of relative pulldown efficiency of GFP-NET1A from lysates of the indicated cell lines with GST-importin α5. n=5. In superplots, data points from individual replicates are color coded, and large, outlined color dots indicate the mean of each replicate. Error bars: SEM. p-values: ***<0.001, ****<0.0001, ns: not significant by Kruskal-Wallis and Kolmogorov-Smirnov test (B) or one-way ANOVA (E-I). Scale bar: 3μm (B), 4μm (D).

The potential for ribosome stalling between the sequences encoding the NLSs and the PH domain raised the possibility of a mechanism whereby the rate of translation elongation could coordinate partner binding by these two targeting domains. To directly test this idea, we aimed to experimentally alter the translation elongation rate within the intervening region encoding the DH domain. To slow down translation, we replaced DH domain-encoding codons with synonymous rare codons. Approximately 20% of codons were replaced leading to a decrease of codon adaptation index from 0.73 to 0.60 (slow mutant). To speed up translation, we mutated the two potential stall sites described above by replacing the proline and one adjacent amino acid with glycine (fast mutant). The presence of slow or fast mutations altered the amount of GFP-NET1A protein produced as expected, i.e. lower amount of protein was produced from the slow mutant, while higher amount was produced from the fast mutant (Figure S7A).

To further validate the impact of these mutations on translation rate, we generated reporter constructs carrying the WT or mutant NET1A coding sequences together with the wild type NET1 5’- and 3’-UTRs (Figure 5B and S7B). 24 copies of the binding site for the MS2 bacteriophage coat protein (MCP) were introduced at the beginning of the 3’UTR to allow visualization of single mRNAs in live cells, through recruitment of multiple copies of an MCP-Halo fusion protein. Furthermore, at the beginning of the coding sequence, a series of SunTag peptide epitopes were introduced, which are recognized by a single chain antibody fragment fused to superfolder-GFP (scFv-GFP), thus allowing detection of new protein chains (Figure 5B and S7B). The wild type, slow or fast variants were expressed under a doxycycline-inducible promoter and scFv signal at translation sites, i.e. in association with individual mRNA molecules, was quantified. This translation signal intensity is proportional to the number of ribosomes bound to any given mRNA. Given equivalent initiation rates, slower or faster elongation would be expected to respectively increase or decrease translation signal intensity (Figure S7B) (Goldman et al., 2021). Indeed, compared to the wild type, the fast mutant exhibits overall lower translation signal intensities, while the slow mutant exhibits the opposite trend (Figure 5B). Taken together, the above data indicate that, for the *NET1* mRNA, translation elongation is limiting and that the introduced mutations alter elongation rates in opposite ways.

### The rate of translation elongation determines partner selection and NET1 protein targeting

We generated stable cell lines that express GFP-NET1A variants (WT, slow and fast) from constructs that carry either the WT or the ΔGA 3’UTR (Figure 5C). The presence of coding sequence mutations did not alter the localization of the expressed mRNAs. Localization was determined by the 3’UTR sequence, such that all mRNAs carrying the WT 3’UTR were distributed peripherally, while mRNAs carrying the ΔGA 3’UTR were perinuclear (Figure 5D, E).

We then assessed how the introduction of these mutations affected binding of GFP-NET1A to its partners and nucleo-cytoplasmic partitioning. Strikingly, slow translation, regardless of mRNA location, led to high interaction with importin and nuclear targeting (Figure 5F, G), and reduced binding to CASK (Figure 5H). In contrast, increasing the rate of translation had the opposite effects, decreasing importin binding and nuclear targeting, while promoting binding to CASK (Figure 5F-H). These data thus show that the location of the *NET1* mRNA and the rate of its translation act in concert to specify NET1 protein interactions and targeting. Perinuclear mRNA and/or slower translation favor NET1-importin interaction and nuclear import, while peripheral mRNA and/or faster translation promote NET1-CASK association and cytoplasmic retention.

As an alternative method to slow down translation we treated cells briefly (30min) with sub-inhibitory concentrations of anisomycin. Under these conditions a subset of ribosomes is expected to be stochastically inhibited, effectively mimicking ribosome stalling. Of note, the concentrations we use here (0.01-0.1 mg/L) are below those leading to peak eIF2α phosphorylation, which is likely induced through a stress response due to ribosome collisions (Figure S8A; (Juszkiewicz et al., 2018; Wu et al., 2020)). In agreement with the above results, we find that slowing translation with sub-inhibitory concentrations of anisomycin leads to increased interaction of peripherally translated NET1A with importin β1 (Figure S8B).

The impact of translation rate suggested that the kinetics and thus likely the order of appearance of NET1 domains could be important. To further test this idea, we generated variants of NET1A in which the NLSs were moved to the C-terminus, i.e. after, rather than before, the PH domain. Interestingly, this placement led to low NET1-importin β1 interaction, even for NET1 translated perinuclearly, and abolished the effect of the mRNA location (Figure S9A, B). These results therefore further support the notion that kinetic competition underlies NET1A partner selection.

It was still possible, however, that the location, rate, or order of protein synthesis alters NET1A-importin interaction by affecting the inherent accessibility of the NLSs, for example through local modifications or altered folding. To assess this, we tested the ability of the various NET1A protein variants within cell lysates to interact with importin *in vitro*. We first identified importin α5/KNPA1 as the importin α member that binds more efficiently to NET1A NLSs (Figure S10) and then tested for its ability to associate with the various NET1A variants in *in vitro* pulldown assays. Interestingly, the increased, or decreased, importin interaction observed *in vivo* was not reflected in corresponding changes of NET1A-importin binding *in vitro* (compare Figure 5F to 5I, and Figure S9B to S9C). Therefore, the impact of the site, rate and order of translation does not appear to be due to alterations in the inherent ability of NET1A variants to recognize importins. Our overall data rather indicate that, *in vivo,* the specific local microenvironment encountered by the nascent, or newly synthesized, polypeptide (likely presenting different concentrations of binding partners) together with the rate at which competing domains appear, favor certain interactions over others and specify the eventual partner selection and target destination.

### mRNA location affects NET1 protein function

Cytoplasmic NET1A is involved in activation of the RhoA GTPase and controls cell adhesion and migration. According to the mechanism described above, the subcellular location of the *NET1* mRNA modulates the nucleo-cytoplasmic distribution of NET1A protein and would thus be expected to influence its function in either compartment. To assess the functional consequences of endogenous *NET1* mRNA localization, we used antisense PMOs to alter its predominantly peripheral distribution to a perinuclear one (Figure 1A). To assess RhoA GTPase activity, we used a FRET-based RhoA activity biosensor (Fritz et al., 2013) and fluorescence lifetime imaging (FLIM) (Figure 6A and S11). Treatment with NET1 PMOs resulted in a reduction in FRET efficiency which was more pronounced in peripheral protrusive regions (Figure 6A). Therefore, perinuclear *NET1* mRNA localization suppresses the cytosolic NET1 function towards RhoA activation, consistent with the observed increased NET1 import into the nucleus. RhoA is a central regulator of the cytoskeleton that influences focal adhesion assembly and maturation. In line with the observed reduced RhoA activity, treatment with NET1 PMOs also led to a significant reduction in the size of paxillin-containing focal adhesions, indicative of a reduced degree of maturation (Figure 6B).

**Figure 6.**
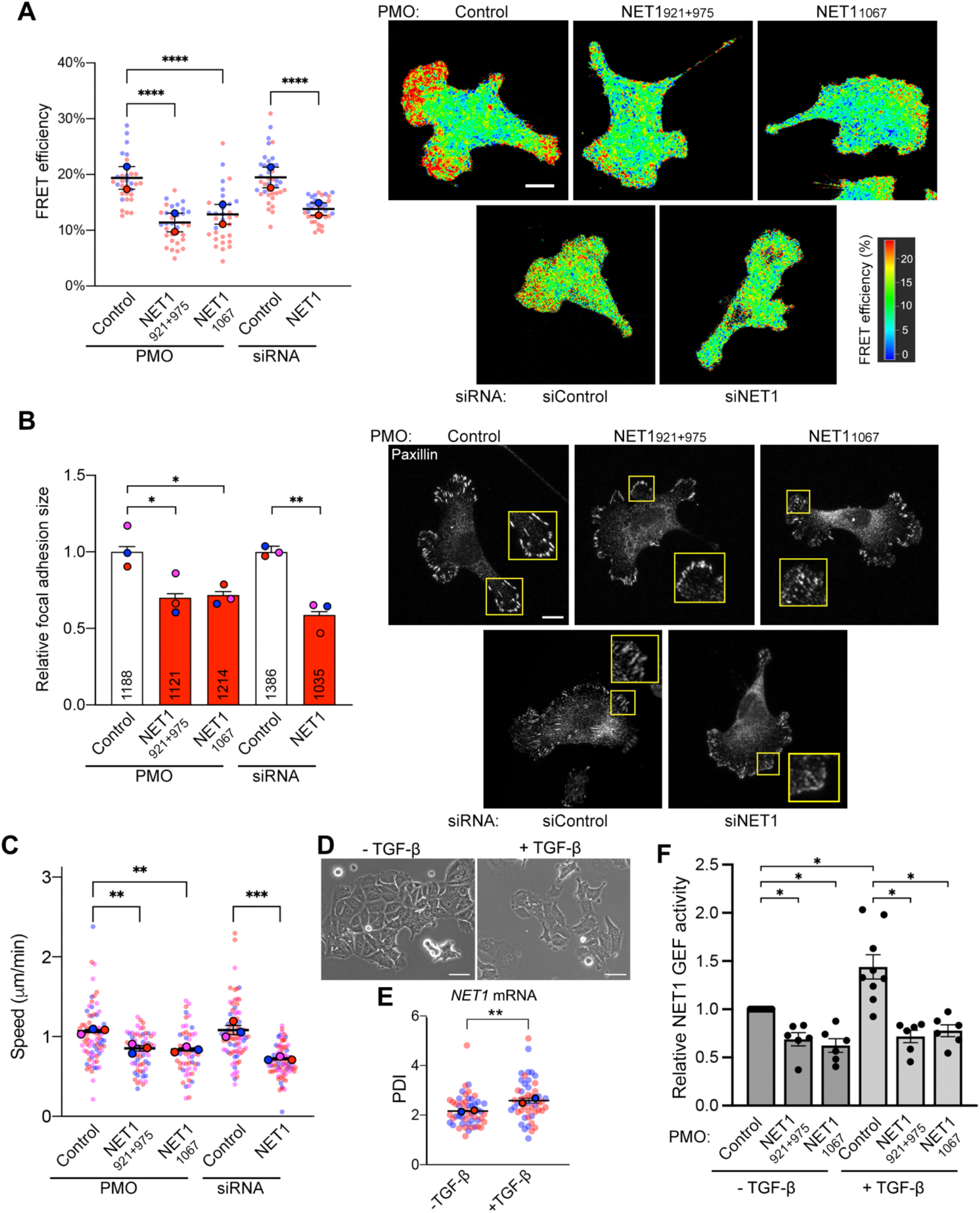
*NET1* mRNA location robustly affects NET1 function in cell migration, and controls TGFβ-induced NET1 activity. **(A)** FLIM imaging of a RhoA activity biosensor in MDA-MB-231 cells treated with the indicated PMOs or siRNAs. Graph shows FRET efficiency in protrusive regions. N=32-38 cells from 2 independent experiments. **(B)** Measurement of focal adhesion size based on paxillin immunofluorescence staining of cells treated with the indicated PMOs or siRNAs. Numbers within each bar indicate the number of focal adhesions measured in 3 independent experiments. **(C)** Average migration speeds measured by time lapse imaging of mCherry-NLS expressing cells treated with the indicated PMOs or siRNAs. N=57-78 cells from 3 independent experiments. **(D)** Phase contrast images of MCF7 cells with or without TGF-β treatment. **(E)** PDI of *NET1* mRNA in MCF7 cells with and without TGF-β. n=50 cells from two independent experiments. **(F)** Relative NET1 binding to nucleotide-free RhoA (RhoA G17A), as a measure of GEF activity, in MCF7 cells with and without TGF-β, and treated with the indicated PMOs. n=3. In superplots, data points from individual replicates are color coded, and large, outlined color dots indicate the mean of each replicate. Error bars: SEM. p-values: *<0.05, **<0.01, ***<0.001, ****<0.0001 by one-way ANOVA. Scale bars: 15μm (A, B); 50μm (D).

To determine whether this alteration in NET1 function was sufficient to cause detectable effects in cell motility, we used cells stably expressing GFP-Lifeact and assessed the speed of protrusive or retracting dynamics of the cell periphery over 1 min intervals (Figure S12A). We further tracked individual cells over long-term time lapse imaging and measured the average cell migration speed (Figure 6C). In both assays, treatment with NET1 PMOs significantly reduced peripheral dynamics and overall cell migration speed, suggesting that peripheral *NET1* mRNA localization is important for efficient cell movement.

### NET1 mRNA location is altered by physiological stimuli

The potential of modulating RhoA activity and cell migration through altering *NET1* mRNA localization, raised the possibility that this mechanism could be utilized under physiological circumstances. We focused on epithelial-mesenchymal transition (EMT) induced by TGF-β, which requires RhoA signaling (Bhowmick et al., 2001). We utilized MCF7 breast epithelial cells, which are known to undergo EMT upon treatment with TGF-β. Indeed, TGF-β induces dissolution and scattering of tightly adherent MCF7 colonies (Figure 6D). Interestingly,

TGF-β also significantly increased the peripheral localization of the *NET1* mRNA (Figure 6E) as well as the amount of NET1 that can bind to nucleotide-free RhoA, indicative of elevated GEF activity (Figure 6F). Importantly, antisense NET1 PMOs that prevent peripheral *NET1* mRNA localization also suppress the TGF-β-induced NET1 GEF activity. Therefore, *NET1* mRNA distribution is altered by a physiological stimulus and is responsible for functional changes linked to RhoA signaling, supporting the biological relevance of the described mechanism.

### NET1 mRNA localization robustly affects NET1 protein functions

To determine the extent to which this mRNA location-dependent mechanism can control NET1 function, we compared the phenotypic effects observed upon forcing a perinuclear *NET1* mRNA distribution to the effects observed upon an almost complete NET1 protein knockdown (Figure S12B). Strikingly, all measured parameters (RhoA activity, focal adhesion size, peripheral dynamics and migration speed) showed a reduction of similar magnitude when NET1 was produced perinuclearly as when its expression was knocked down (Figure 6A-C and S12A). Therefore, even though perinuclearly translated NET1 is expressed at normal levels (Figure 1E and S2E,F), its efficient nuclear import strongly suppresses its ability to function in the cytoplasm, phenocopying acute NET1 protein loss. We conclude that altering *NET1* mRNA distribution does not simply provide a subtle regulatory mechanism but can robustly influence partner selection and consequently the targeting of the protein and its function between its two subcellular destinations.

## Discussion

Numerous proteins are targeted to multiple subcellular destinations where they can exert distinct moonlighting functions or promote diverse functional outcomes (Avadhani, 2011; Singh and Bhalla, 2020). The mechanisms specifying the balance between distinct functions or destinations are not always clear but are largely thought to involve post-translational events. For example, the distribution of NET1 between the nucleus and the cytoplasm can be controlled post-translationally through protein modification (Song *et al*., 2015; Ulu *et al*., 2018). We show here that a distinct level of control operates during, or early after, protein synthesis and plays a major role in determining the functional potential of the protein. Specifically, the location of the mRNA in the cytoplasm and the rate of its translation act in combination to specify the targeting, and thereby the functional potential, of the encoded NET1 protein. NET1 targeting between the nucleus and cytoplasm is determined through antagonistic binding of importins or CASK to the NLS and PH domain respectively. Our model (Figure 7) suggests that partner selection is determined by kinetic competition and is influenced either by the rate of appearance of the interacting domains or by presenting the newly synthesized protein into different local environments that likely differ in the concentration of binding partners. In this way, *NET1* mRNA translation at the periphery disfavors NLS-importin interaction and instead promotes PH domain-CASK binding, thus retaining the protein in the cytoplasm. This balance can be shifted either by moving the mRNA to a perinuclear location, where the NLSs are more efficiently recognized by the higher importin concentration, or by slowing down translation thereby allowing more time for NLS-importin interaction.

**Figure 7:**
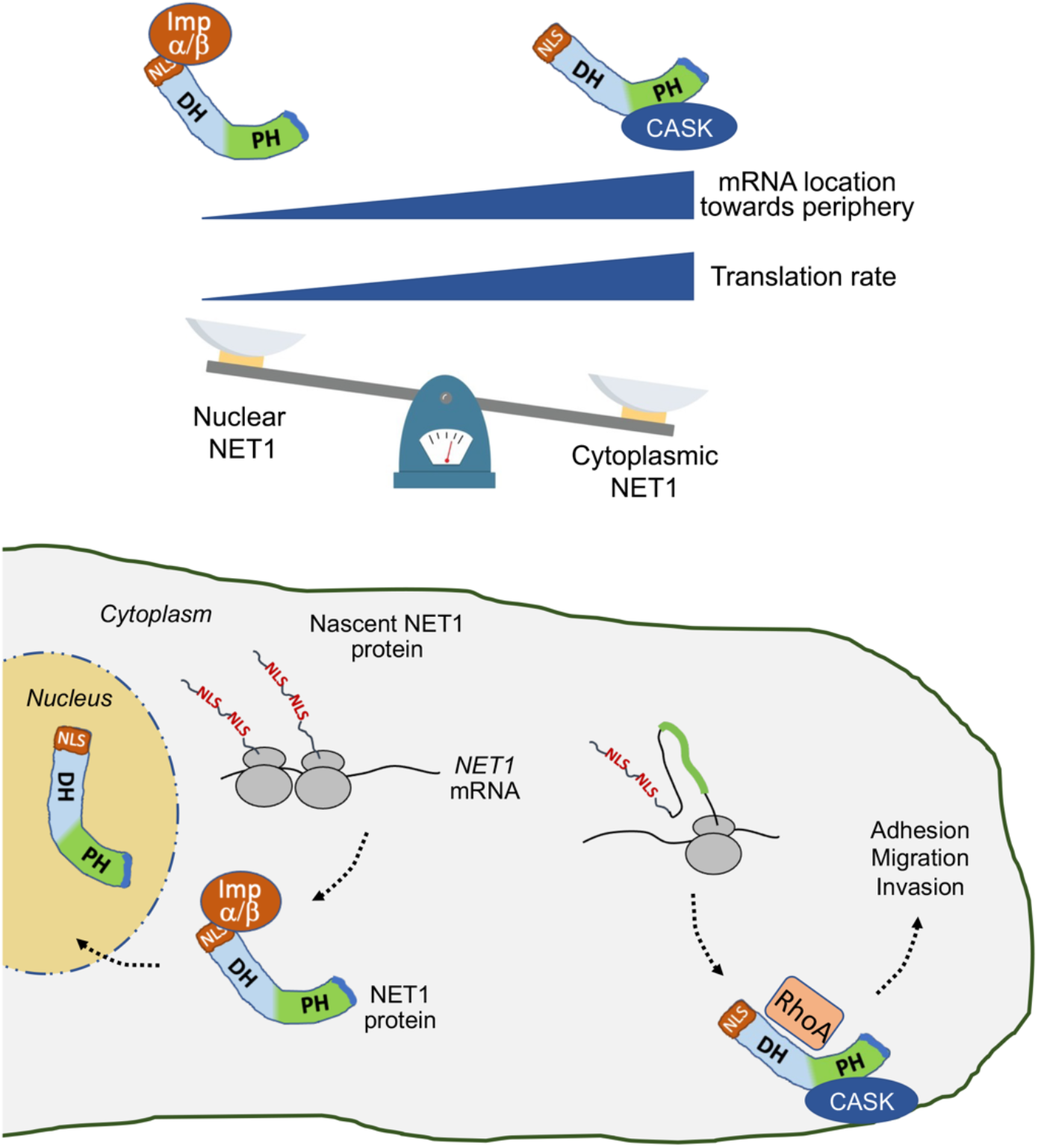
Proposed model. Binding of NET1 to importins or to CASK, and NET1 targeting to the nucleus or cytoplasm, is determined by the location of the *NET1* mRNA and the rate of its translation. Perinuclear mRNA and/or slower translation rate favor interaction of the N-terminal NLSs with import receptors and targeting to the nucleus. Peripheral mRNA and/or faster translation rate favor CASK binding through the PH domain to competitively retain NET1 in the cytoplasm. Cytoplasmic NET1 can activate RhoA and promote cell adhesion and migration.

Our data indicate that, in the case of NET1A, a choice for eventual protein targeting occurs during or shortly after translation. It is well appreciated that protein folding and association with partners can occur as soon as the nascent protein chain emerges from the ribosome and such co-translational events can ensure proper protein folding and prevent promiscuous and non-productive interactions (Kramer et al., 2019). Co-translational events can be influenced by the rate of translation and by ribosome pausing to ensure proper complex assembly (Collart and Weiss, 2020). They have also been proposed to play a role towards directing interactions of proteins when multiple potential functional options are possible (Schwarz and Beck, 2019). It is quite intuitive that the potential co-translational associations that a nascent chain can engage in will be shaped by the immediate microenvironment in which protein synthesis takes place. Supporting this idea, local co-translational interactions, determined by mRNA location, have been shown to direct the functional potential of signaling proteins (Moissoglu *et al*., 2020). The evidence presented here lends experimental support to the idea that co-translational events serve not only a quality control role, but in combination with mRNA targeting can be used as a selection mechanism for proteins that engage in multiple potential functional complexes.

Such a partner selection mechanism based on mRNA targeting could be widely utilized. A large fraction of cellular mRNAs are not randomly distributed but adopt complex and diverse distribution patterns in the cytoplasm. While the functional roles served by these targeting events have been studied in only a handful of cases, large-scale correlations offer some interesting insights. Proteins expressed by mRNAs localized to distal locations tend to have a larger number of interaction partners compared to proteins expressed from non-targeted mRNAs (Weatheritt et al., 2014). Exclusively mitochondrial proteins are more likely to be expressed from mRNAs that are targeted to mitochondria compared to mitochondrial proteins that exhibit dual localization (Ben-Menachem et al., 2011). We suggest that mRNA localization and the kinetics of co-translational events, driven by translation rate and the local concentrations of binding partners, can provide a widely-used mechanism for partner selection and thus for specifying protein distribution and function.

Our data cannot currently discern whether mRNA location and translation rate are independently modulated or whether they are coupled. Understanding if an mRNA is translated with different rates at different cellular locations will be an intriguing area for future investigation.

## Acknowledgments

We thank the CCR Genomics Core of the National Cancer Institute, NIH for ddPCR analysis. This work was funded by the Intramural Research Program of the Center for Cancer Research, National Cancer Institute (NCI), National Institutes of Health (NIH) (1ZIA BC011501 to S. Mili); by the Intramural Research Program of the National Institute of Diabetes and Digestive and Kidney Diseases (NIDDK), NIH (DK075132 to N.R.G.); and by the Postdoctoral Research Associate Training Program (PRAT) at the National Institute of General Medical Sciences (NIGMS) (1FI2GM137845 to S. Meydan).

## Author Contributions

S. Mili, A.N.G. and K.M. conceived the project and designed experiments. A.N.G. and K.M. performed experiments. A.N.G., K.M. and S. Mili analyzed data. S.P. provided image analysis code. S. Meydan and N.R.G. performed and analyzed ribosome profiling. S. Mili and A.N.G. wrote original manuscript. All authors reviewed and edited the manuscript.

## Methods

### Cell culture and generation of stable cell lines

MDA-MB-231 cells were obtained from ATCC (cat # HTB-26) and cultured in Leibovitz’s L-15 medium (Gibco) supplemented with 10% FBS and 1% Penicillin-Streptomycin at 37°C in atmospheric air in a humidified cell culture incubator. MCF7 (obtained from ATCC; cat# HTB-22) and 293 TREx cells (ThermoFisher, cat# R780-07) were cultured in Dulbecco’s modified eagle medium (DMEM; Gibco) supplemented with 10% FBS and 1% Penicillin-Streptomycin at 37°C and 5% CO_2_. NIH/3T3 mouse fibroblast cells were cultured in DMEM supplemented with 10% calf serum, sodium pyruvate and 1% Penicillin-Streptomycin at 37°C and 5% CO_2._ Cells were passaged by trypsinization using 0.05% trypsin (Gibco). Cells used in this study have tested negative for mycoplasma.

To generate stable cell lines expressing GFP-tagged NET1A and various mutants, MDA-MB-231 cells were infected with the corresponding lentiviruses and selected with 6μg/mL blasticidin (ThermoFisher). Cells were sorted by FACS to select for a low level of expression.

To generate stable cell lines expressing the doxycycline-inducible RhoA FRET biosensor, MDA-MB-231 cells were infected with the corresponding lentivirus and selected with 0.6mg/mL geneticin (ThermoFisher). To prevent leaky expression of the biosensor, these cells were maintained in media supplemented with 10% Tet-system approved FBS (ThermoFisher). To induce RhoA FRET biosensor expression, cells were treated overnight with 1μg/mL doxycycline.

To generate stable cell lines expressing FLAG-KPNB1, LifeAct-GFP, or Cherry-NLS, MDA-MB-231 cells were infected with the corresponding lentiviruses and selected with 1µg/mL puromycin (ThermoFisher) or fluorescent cells were sorted by FACS.

To generate stable cell lines expressing translation reporters, NIH/3T3 cells were sequentially infected with lentiviruses expressing stdMCP-stdHalo (Addgene #104999, modified to remove the Kozak sequence and ATG initiating codon from the sequence of the first HaloTag) and scFv-GFP (derived from Addgene #60906 after introduction of a stop codon before the NLS sequence). A uniformly expressing cell population was selected by fluorescence activated cell sorting. Reporter constructs expressing NET1A variants (WT, slow or fast) were then introduced through lentiviral infection, and stably expressing lines were selected with Geneticin (Thermo Fisher Scientific). Expression of the reporters was induced by addition of 1 μg/ml doxycycline approximately 3-4 hrs before imaging.

For protein synthesis inhibitor studies, cells were incubated with either cycloheximide (100µg/mL) or puromycin (100µg/mL) for 20 min at 37°C. For TGF-β treatment, cells were plated in 1% FBS-containing DMEM for 72 hrs with or without 4ng/mL human recombinant TGF-β (Peprotech; cat #100-21).

### Plasmid constructs and lentivirus production

To express N-terminally GFP-tagged NET1A with different UTRs, the coding sequence of EGFP was ligated to the coding sequence of human NET1A. This fusion was then ligated to fragments corresponding to the 3’UTR of human NET1 (NM_001047160.3; wt UTR) or the 3’UTR of NET1 that is missing 78 nucleotides (921-999) (ΔGA UTR). These fragments were then cloned into the pCDH-PGK lentiviral vector. To generate NET1A deletion mutants, the coding sequence of human NET1A, lacking the DH or PH domain, or the PDZ motif, was synthesized as a gBlock gene fragment (Integrated DNA Technologies). Each fragment was ligated to an N-terminal GFP-tag and to the wt NET1 3’UTR into the pCDH-PGK lentiviral vector. To generate the NET1A with mutated NLSs or the slow or fast mutants, gBlock gene fragments (Integrated DNA Technologies) carrying the corresponding mutations were synthesized. To generate NET1A with C-terminal NLSs, the N-terminal NET1A region encompassing the NLSs was PCR amplified from the gBlock fragment and ligated in frame at the C-terminus of a fragment containing the DH-PH domains and PDZ motif of NET1A. Each fragment was ligated to an N-terminal GFP-tag and to the wt or ΔGA NET1 3’UTR into the pCDH-PGK lentiviral vector. See Supplementary Table S1 for exact sequence information of fragments used.

To generate the NET1 translation reporter cell lines the 5’UTR sequence of human NET1A was synthesized as a gBlock gene fragment (Integrated DNA Technologies). A fragment containing 24xGCN4_v4 repeats was derived from Addgene plasmid #74928. Those were cloned upstream of gBlock gene fragments containing the human wild type NET1A coding sequence or slow and fast mutant variants. A fragment with 24xMS2v7 repeats was obtained from Addgene plasmid #140705 and cloned downstream of the coding sequence, followed by the wild type human NET1 3’UTR. Fragments were cloned into the pInducer 20 lentiviral vector (Addgene plasmid #44012) for doxycycline-inducible expression.

To generate the RhoA FRET biosensor, the RhoA biosensor fragment was PCR amplified from the pLentiRhoA2G plasmid (Addgene #40179) to add BamHI and XhoI sites at the ends. Site directed mutagenesis was used to introduce either a T19N (dominant negative) or Q63L (constitutively active) mutation. The wild type or mutant fragments were cloned into the pENTR- 1A vector and a recombination reaction using the Gateway LR Clonase II enzyme mix (ThermoFisher) was performed to move the RhoA FRET sensor fragment into the doxycycline-inducible pInducer20 vector (Addgene plasmid # 44012).

To express FLAG-tagged importin β (KPNB1), the KPNB1 coding sequence was PCR amplified from the KPNB1 ORF clone plasmid (Origene, cat# RC200659) and XbaI and BamHI restriction sites were added to the ends of the fragment. This fragment was cloned into the pCDH-CMV-EF1-MCS-Puro lentiviral vector.

To express FLAG-tagged CASK, a lentiviral plasmid that contained the human CASK CDS (NM_001367721.1) with an N-terminal FLAG tag driven by the human PGK promoter with a puromycin resistance gene driven by the CMV promoter was purchased from VectorBuilder. The mCherry-FLAG-tagged CASK construct (VectorBuilder) has mCherry at the C-terminus of the FLAG-CASK CDS.

Lentiviruses were produced in HEK293T cells. The cells were transfected with the lentiviral vectors together with the pMD2.G and psPAX2 packaging plasmids using PolyJet In Vitro DNA Transfection Reagent (SignaGen) for 48 hrs. Harvested virus was precipitated with polyethylene glycol overnight at 4°C.

### Morpholino and siRNA transfection

For knockdown experiments, 40 pmoles of siRNA were transfected into cells by Lipofectamine RNAiMAX (ThermoFisher, cat# 13778-150) according to the manufacturer’s protocol. Cells were assayed 72 hours post-transfection. The following siRNAs were used: AllStars negative control (Qiagen cat# 1027281) and siNET1 #4 (Qiagen cat# SI00082040; target sequence: 5’- ACGGAAAGAGACTTTGGTGTA-3’), siCASK #5 (Qiagen cat# SI02223368; target sequence: 5’-AACCAATGGGAATCACTTTAA-3’) and siCASK #10 (Qiagen cat# SI04437720; target sequence: 5’-CAGACCGGTTTGCGTACCCTA-3’).

Morpholino oligos (Gene Tools, LLC) used are listed in Supplementary Table S2. Morpholinos were transfected using Endoporter (Gene Tools, LLC). Cells were assayed 72 hours post-transfection.

### Western blot

The following primary antibodies were used: rabbit anti-GFP (1:1000; ThermoFisher cat# A-11122), rabbit anti-NET1 (1:1000, Abcam cat# ab113202), rabbit anti-importin β1 (1:1000, Cell Signaling Technology cat# 60769), mouse anti-CASK (1:1000, Santa Cruz cat# sc-13158), rabbit anti-CASK (1:1000; Invitrogen cat# PA1544), mouse anti-tubulin (1:2000; Sigma cat# T6199), rabbit anti-GAPDH (1:2000; Cell Signaling cat# 2118L). Anti-rabbit and anti-mouse secondary antibodies from Li-Cor were used at 1:10,000. Membranes were scanned using an Odyssey fluorescent scanner (Li-Cor) and bands were quantified using ImageStudioLite (Li-Cor).

### RNA Fluorescence in situ hybridization

**(FISH)** MDA-MB-231 and MCF7 cells were plated on collagen IV-coated (10µg/mL) glass coverslips for 2 hours and fixed with 4% paraformaldehyde for 20 mins at room temperature. FISH was performed using the ViewRNA ISH Cell Assay kit (ThermoFisher, cat# QVC0001) according to the manufacturer’s protocol. The following Affymetrix probes were used in this study: human NET1 (cat # VA6-3169338), human RHOA (cat # VA6-14829-01) and GFP (cat # VF6-16198). Green cell mask (ThermoFisher) was used to identify the cell border and samples were mounted with ProLong Gold antifade with DAPI.

### Focal adhesion immunofluorescence

MDA-MB-231 cells plated on collagen IV-coated (10µg/mL) coverslips were fixed with 4% paraformaldehyde in PBS for 15 mins at room temperature then permeabilized with 0.2% Triton X-100 in PBS for 5 mins at room temperature. Coverslips were blocked for 1 hr in 5% goat serum in PBS for 1 hr at room temperature. Mouse anti-paxillin primary antibody was then added at 1:300 for 1.5 hrs. After washing, Alexa Fluor-488 Phalloidin (1:800; Invitrogen cat# A12379) and anti-mouse Alexa Fluor-647 secondary (1:500; Invitrogen) were added for 1 hr. Samples were mounted in ProLong Antifade mounting media with DAPI.

### Co-Immunoprecipitation

Cells were lysed in an ice-cold buffer containing 50mM Tris pH 7.4, 1% NP-40, 150mM NaCl, 10mM MgCl_2_, 10% glycerol and Halt protease and phosphatase inhibitor cocktail (ThermoFisher). Lysates were cleared by centrifugation at 4°C and added to GFP-Trap Magnetic agarose beads (Chromotek, cat #gtma-10) for 1 hr at 4°C with rotation. After washing with lysis buffer, immobilized complexes were eluted with Laemmli’s buffer by boiling for 5 minutes. The supernatant was separated from the beads by a magnet and then analyzed by SDS-PAGE and immunoblotting.

For the endogenous CASK co-immunoprecipitation experiments, cells were lysed in an ice-cold buffer containing 50mM Tris pH 7.4, 1% TritonX-100, 75mM NaCl, 10mM MgCl_2_, 10% glycerol and Halt protease and phosphatase inhibitor cocktail (ThermoFisher). After 5 minutes of lysis, lysates were sonicated for 10 seconds on intensity setting #2 using a sonicator (Misonix Inc. Sonicator XL). Lysates were gently spun for 5 mins at 4,000 rpm. The supernatant was then incubated as described above for the other co-immunoprecipitation experiments.

### Proximity Ligation Assay (PLA)

Cells plated on collagen IV-coated (10µg/mL) coverslips were washed in PBS and fixed with 4% paraformaldehyde in PBS for 15 min at room temperature then permeabilized in 0.2% Triton X-100 for 5 min at room temperature. The DuoLink In Situ Red Kit (Sigma, cat# DUO92008) was used for PLA and the manufacturer’s protocol was followed. Briefly, cells were blocked in the provided blocking buffer for 1 hr at 37°C in a humidified chamber. Primary antibodies were diluted in the provided DuoLink antibody diluent solution and incubated on the cells for 1.5 hrs at room temperature in a humidified chamber. The following primary antibodies were used: rabbit anti-GFP (1:200; ThermoFisher cat# A11122) and mouse anti-FLAG M2 (1:800; Sigma cat# F1804). After washing, the PLA probes supplied with the kit were used at a 1:10 dilution in the antibody diluent buffer provided and incubated for 1 hr at 37°C. The ligation step was performed for 30 min at 37°C then amplification for 100 min at 37°C. After these steps, the coverslips were washed and fixed with 4% paraformaldehyde in PBS for 10 min at room temperature then stained with Alexa Fluor 488 Phalloidin (1:500; Invitrogen cat# A12379) in blocking buffer for 20 min at room temperature and mounted in DuoLink PLA mounting medium with DAPI. Slides were kept at 4°C in the dark until imaging the next day.

### Active NET1 pull down assay

MCF7 cells transfected with control or NET1-targeting morpholinos, with or without TGF-β, were rinsed with ice cold HBS buffer (20mM HEPES pH 7.5, 150mM NaCl) and lysed with ice cold HBS lysis buffer (20mM HEPES pH 7.5, 150mM NaCl, 5mM MgCl_2_, 1% TritonX-100) with Halt protease and phosphatase inhibitor for 10 mins then centrifuged at 4°C to clear the lysate. Cleared lysate was added to a GST-RhoA(G17A) bead slurry for 1 hr at 4°C with rotation. The beads were washed x5 with cold HBS lysis buffer and proteins were eluted with Laemmli’s buffer by boiling for 5 minutes before running on an SDS-PAGE gel.

### In vitro pull down assay

Cells stably expressing GFP-NET1A variants were rinsed with ice cold PBS and lysed in 1mL of HEGMN buffer (25mM HEPES, 100mM KCl, 12.5mM MgCl_2_, 0.1mM EDTA, 10% glycerol, 0.1% NP-40) supplemented with Halt phosphatase and protease inhibitor on ice for 5 mins. For pull down, 100uL of Pierce glutathione magnetic agarose beads (ThermoFisher, cat#78601) were incubated with the lysates and with 1.0µg, or with the indicated amounts, of GST-KPNA recombinant human protein (Abnova, GST-KPNA1 (cat#H00003836-P01), GST-KPNA2 (cat#H00003838-P01), GST-KPNA4 (cat#H00003840-P01)) for 1 hour with rotation at room temperature. Beads were washed x5 with lysis buffer, proteins were eluted with 35μl reducing buffer, boiled for 5 mins, and analyzed by Western blot.

### Microscopy and Image Analysis

Focal adhesion size, RNA FISH and PLA experiments were imaged using a Leica SP8 confocal microscope with an HC PL APO 63x oil immersion objective. Z-stacks through the cell volume were obtained and maximum intensity projections were used for all analysis.

For FISH images, calculation of the PDI index was performed using a previously published custom Matlab script(Stueland *et al*., 2019).

For GFP-NET1A/FLAG-importin PLA experiments, an ImageJ script based on the Analyze Particles plugin was used to calculate the number of PLA dots inside each cell. Equal thresholding among all images was used during analysis. For GFP-NET1A/FLAG-CASK PLA, a series of custom ImageJ macros were used to define an inner boundary 3μm from the cell edge and to obtain PLA signal intensity within this 3μm peripheral zone, after background subtraction.

For live cell imaging of GFP-NET1A nuclear and cytoplasmic ratios, cells were plated on glass-bottom dishes and DRAQ5 (ThermoFisher) was added to the cell culture media at a concentration of 1µM for 30 minutes prior to imaging. Cells were placed inside an incubation chamber mounted on the microscope stage to keep cells at 37°C. Z-stack images through the cell volume were taken and imported into Imaris (Bitplane AG) where DRAQ5 staining was used to determine the nuclear volume. Surfaces were created within the software to segment out the nuclear region from the whole cell and the GFP intensities were calculated for both regions.

For cell migration experiments, an Olympus IX81 inverted microscope was used with an incubation chamber around the microscope to keep the cells at 37°C for the duration of the experiment. To facilitate tracking, cells expressing Cherry-NLS were used. A 10x objective was used for imaging and time lapse images were taken at 10 min intervals for a duration of 8 hours. Images were analyzed in ImageJ software using the Manual Tracking plugin. The raw tracking coordinates were imported into the DiPer macro(Gorelik and Gautreau, 2014) for cell migration speed analysis in Microsoft Excel.

To analyze the size of focal adhesions, images were converted to binary masks in ImageJ by thresholding and the analyze particles plugin was used to measure focal adhesion size. Equal thresholding among all images were used during analysis.

To analyze cellular membrane dynamics, cells expressing LifeAct-GFP were plated on collagen IV-coated glass bottomed dishes and imaged on a heated stage incubator chamber on a Leica SP8 confocal microscope. Time lapse Z-stack images were taken throughout the whole Z-plane of the cell every 1 minute for 1 hr total. To measure the speed of membrane edge protrusion and retraction, the ADAPT ImageJ plugin was used(Barry et al., 2015). The images were thresholded within the plugin until the background outside the cell was subtracted out and the outline of the cell remained. The curvature window setting was kept at the default of 10.

To measure RhoA FRET activity, cells expressing the biosensor were plated on collagen IV-coated glass bottomed dishes and imaged on a heated stage incubator chamber on a Leica SP8 confocal microscope. The LAS X Falcon FLIM module was used to analyze the data. To measure FRET efficiency in protrusive and non-protrusive areas, an ROI was made to include all the cell’s protrusions and compared to the FRET value of the donor-only expressing cells in a phasor plot. To measure FRET efficiency in non-protrusive areas, an ROI box was made along the cell membrane in an area without membrane protrusions. This FRET value was compared to the donor-only cell FRET value in a phasor plot.

Live imaging of cells expressing translation reporters was done using a Nikon Eclipse Ti2-E inverted microscope, equipped with a motorized stage, a Yokogawa CSU-X1 spinning disk confocal scanner unit, and operated using NIS-elements software. Acquisitions were performed using an Apochromat TIRF 100× oil immersion objective (N.A. 1.49, W.D. 0.12 mm, F.O.V. 22 mm) and Hamamatsu ORCA-Fusion BT Gen III back-illuminated sCMOS cameras. Constant 37°C temperature and 5% CO_2_ were maintained using a Tokai Hit incubation system. To label MCP-Halo proteins, cells were supplemented with 200nM of JFX554 HaloTag ligand, obtained from Janelia Research Campus for 2 hrs. The medium was then replaced, and 1 μg/ml doxycycline was added to induce expression of reporter mRNAs for 3 hrs. Cells were plated on fibronectin (5 mg/mL)-coated 35 mm glass bottom dishes for ∼1 hr, and samples were simultaneously excited using a 488 nm and 561nm laser lines. Emissions were recorded on two separate cameras, which were aligned prior to every imaging session using fluorescent TetraSpeck beads (0.1, 0.5 and 4 μm in diameter). Individual mRNA and translation particles were identified using the TrackMate plugin in ImageJ/Fiji, and a custom Matlab script was used to assign translation spots to individual mRNA particles (within a 400nm radius) and correct for cell-specific background.

### Ribo-seq and Disome-seq experiments

HEK293 cells grown to a confluency of ∼80% in a T-75 flask were washed with ice-cold DPBS and flask frozen in liquid nitrogen. 400 μL of lysis buffer (20 mM Tris-HCl pH 7.5, 150 mM NaCl, 5 mM MgCl_2_, 1 mM DTT, 100 μg/mL cycloheximide, 1% Triton X-100, 25 U/mL Turbo DNase) was added to the frozen cells and the cells in lysis buffer were thawed on ice. Cells were scraped from the flasks and lysates were transferred to Eppendorf tubes, followed by incubation on ice for 10 min. After passing the lysates through 25 G needles ten times, lysates were clarified by centrifugation for 10 minutes at 21,000 g at 4°C and supernatants were kept for library preparation. The total RNA concentration in the lysates were measured by using Quant-it RiboGreen RNA Assay Kit (ThermoFisher, cat#R11490) and the lysate containing 30 μg of RNA was digested with 60 U of RNase I (ThermoFisher, cat#AM2295) for 1 hour at room temperature to capture both monosome and disome footprints as previously described(Mito et al., 2020). RNase I digestion was stopped by the addition of 10 μL Superase-In (ThermoFisher, cat#AM2694) and lysates were centrifuged for at 21,000 g at 4°C. Supernatants were loaded onto a 1 M sucrose cushion and then pelleted by ultracentrifugation by using a TLA100.3 rotor (Beckman Coulter, cat#349490) for 1 hour at 100,000 rpm at 4°C. The ribosome pellet was resuspended in 700 μL TRIzol (ThermoFisher, cat#15596026) and RNA was extracted as recommended by the manufacturer. Concentration of the RNA was determined by a NanoDrop UV spectrophotometer and 2 μg of RNA was loaded onto 15% Criterion-TBE-Urea polyacrylamide gel (Biorad, cat#3450093) for size selection. For Ribo-seq and Disome-seq libraries, gel regions corresponding to 25-34 nucleotides and 50-70 nucleotides, respectively, were cut. RNA was eluted overnight from the gel slice (0.3 M sodium acetate, 1 mM EDTA and 0.25 % SDS) and isopropanol precipitated the next day. The extracted footprints were dephosphorylated and pre-adenylated linkers were ligated as described previously(McGlincy and Ingolia, 2017). rRNAs were removed by Qiagen FastSelect kit (cat#334386) during reverse transcription step as described by the manufacturer. Quality of the final libraries were assessed by Bioanalyzer 2100 (Agilent, cat#G2939BA) using the High Sensitivity DNA kit (Agilent, cat#5067-4626). Sequencing was performed on a NovaSeq sequencer at the NIH/NHLBI DNA Sequencing and Genomics Core.

### Processing of Ribo-seq/Disome-seq data

Sequencing data was processed as described previously(Karasik et al., 2021). Briefly, the fastq files of Ribo-seq and Disome-seq data were trimmed to remove linkers and demultiplexed to isolate individual samples from pooled data using Cutadapt. Contaminating tRNAs and rRNAs were filtered out by Bowtie1 allowing two mismatches in -v mode and -y option to increase sensitivity of the alignment. The noncoding RNA fasta file was created by downloading rRNA sequences from the SILVA project (release 128)(Quast et al., 2013) and tRNA sequences from GtRNAdb (H. sapiens release 16)(Chan and Lowe, 2009). After filtering out the non-coding RNAs, a custom python script was used to remove PCR duplicates by comparing the 7-nucleotide unique molecular identifiers (UMIs) in libraries. The UMIs were then trimmed by Cutadapt and reads were aligned to a transcriptome with bowtie1 using the parameters -v 1 (one mismatch allowed), -y and -p 12. The transcriptome, RefSeq Select+MANE (ncbiRefSeqSelect) was downloaded from UCSC on April 14, 2020 and used for alignment after removal of duplicates on alternative chromosomes. The footprint length distribution was obtained by using FastQC, version 0.11.7 (Babraham Bioinformatics). For Ribo-seq and Disome-seq data, only the reads between 25-34 and 58-65 nucleotides were analyzed, respectively.

We used custom python3 scripts (https://github.com/guydoshlab/ribofootPrinter)(Guydosh, 2021) to create files with normalized mapped reads and used writegene2 script to visualize ribosome and disome reads mapped to the *NET1* gene.

### Statistical analysis

All statistical analysis was performed using GraphPad Prism software using the appropriate statistical tests as indicated within the text and figure legends. Normally distributed datasets were analyzed using parametric statistical tests. Datasets deviating from a normal distribution were analyzed using non-parametric tests. Follow up tests were included, as appropriate, to adjust for multiple comparisons.

## Data availability and Code availability

All data and analysis codes are available from the corresponding author upon reasonable request.

## Supplementary Tables

**Table S1:** Sequences of coding region fragments used for generation of plasmid constructs

**Table S2:** Sequences of morpholino oligonucleotides (PMOs) used in the study

---

**Figure S1:**
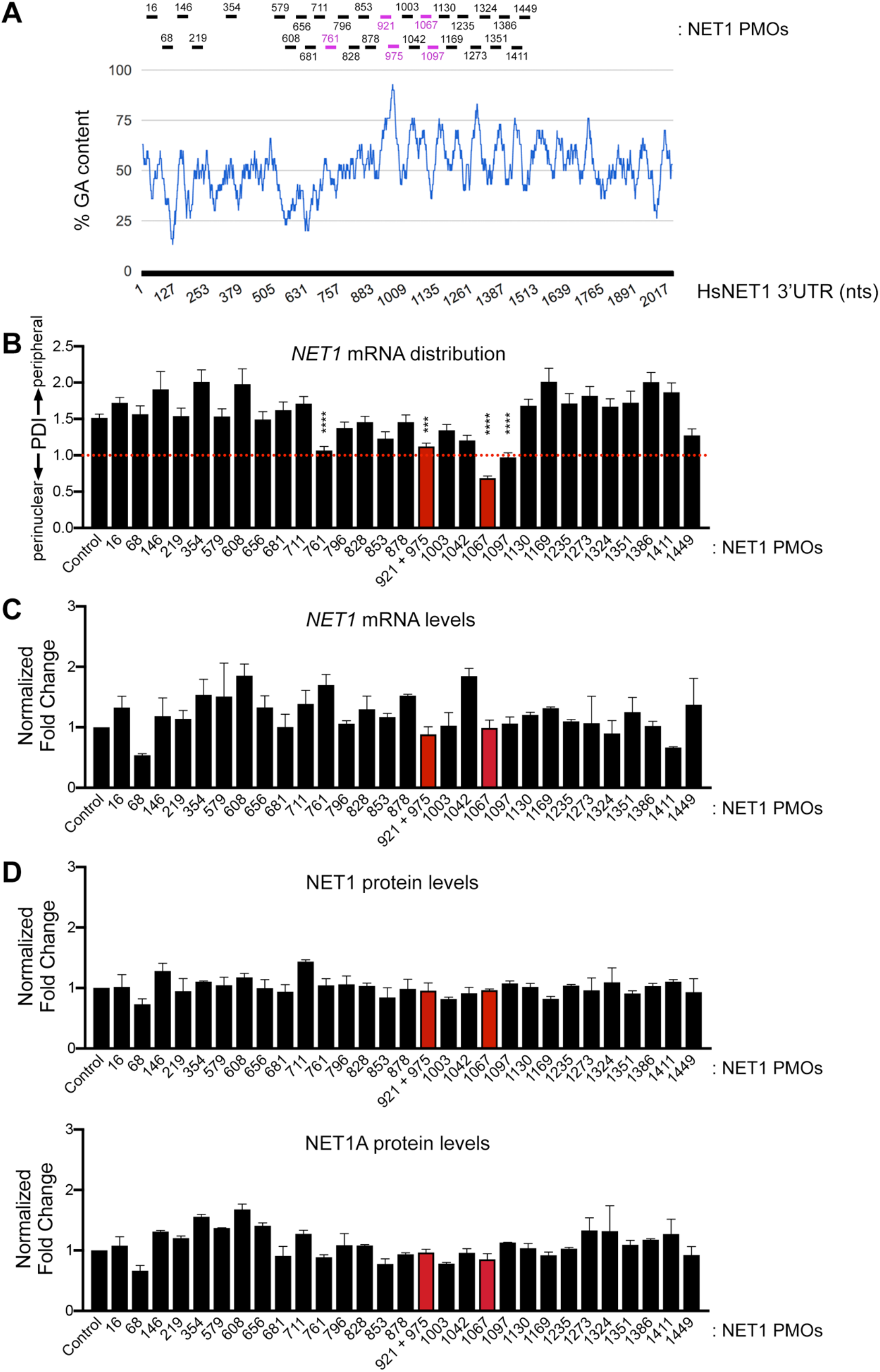
Antisense oligonucleotide tiling across the 3’ UTR of the *NET1* mRNA. **(A)** Chart showing the percent GA nucleotide content within the 3’ UTR of the human *NET1* mRNA. The NET1 PMOs tested are shown above the chart and the PMOs in magenta are those that showed significant reduction in PDI (see B). **(B)** PDI quantification of *NET1* mRNA distribution determined by FISH of cells treated with the indicated PMOs. n=30-118 cells. PDI=1 indicates a diffuse distribution, PDI>1 indicates peripheral distribution, PDI<1 indicates a perinuclear distribution. **(C)** *NET1* mRNA levels determined by ddPCR after treatment with the indicated PMOs. n=2. **(D)** Protein levels of the NET1 and NET1A isoforms, by Western blot of cells treated with the indicated PMOs. n=2. Error bars: SEM. p-values: ***<0.001, ****<0.0001 by one-way ANOVA. The red bars indicate the *NET1* mislocalizing oligonucleotides used throughout this study.

**Figure S2:**
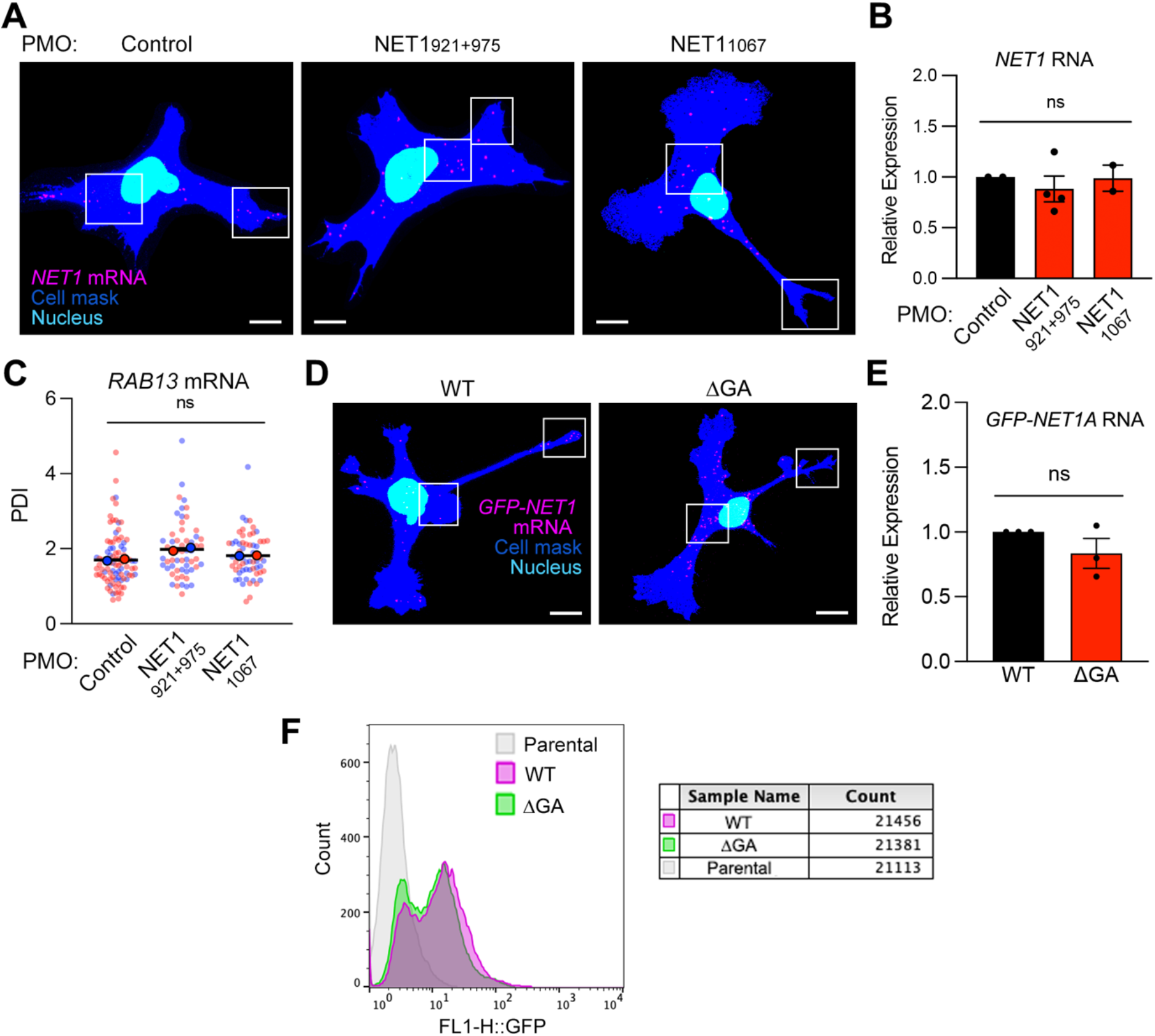
Specificity and effect of altering *NET1* mRNA localization on mRNA levels and protein expression. **(A)** Representative RNA FISH images of MDA-MB-231 cells treated with the indicated PMOs. Boxed regions indicate areas shown in Figure 1A. **(B)** *NET1* mRNA expression measured by ddPCR of PMO treated cells. n=2-4. **(C)** PDI quantification of *RAB13* mRNA distribution in cells treated with the indicated PMOs. NET1-targeted PMOs do not alter the distribution of the co-regulated *RAB13* mRNA. **(D)** Representative RNA FISH images of cells expressing the WT and ΔGA GFP-NET1A constructs. Boxed regions indicate areas shown in Figure 1D. **(E)** *GFP-NET1A* mRNA expression as measured by ddPCR of the indicated stable cell lines. n=3. **(F)** Flow cytometry analysis of GFP fluorescence in parental MDA-MB-231 cells and GFP-NET1A cells with either the WT or ΔGA UTR. Error bars: SEM. ns: not significant by one-way ANOVA (B, C) or unpaired t-test (E).

**Figure S3:**
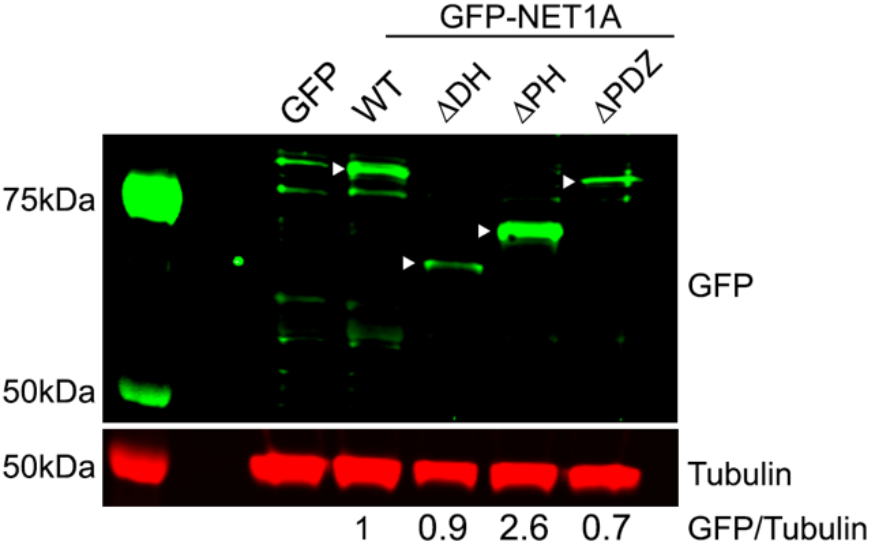
Characterization of GFP-NET1A mutant variants. Western blot of stable cell lines expressing GFP or the indicated GFP-tagged NET1A variants. White arrowheads indicate the band of interest for each lane. Tubulin was used as a loading control and relative expression levels are indicated at the bottom.

**Figure S4:**
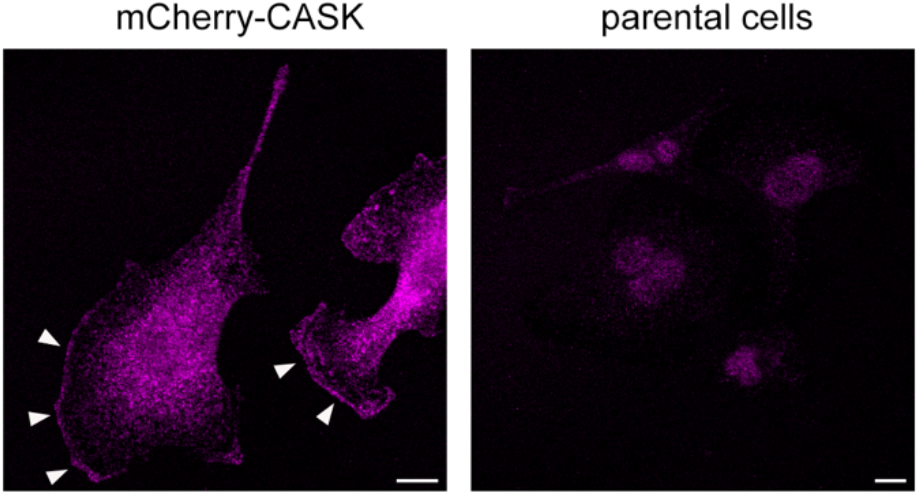
Intracellular CASK distribution. Immunofluorescence of mCherry-CASK expressing cells. Arrowheads indicate peripheral areas of CASK accumulation, potentially reflecting sites of CASK association with membranes.

**Figure S5:**
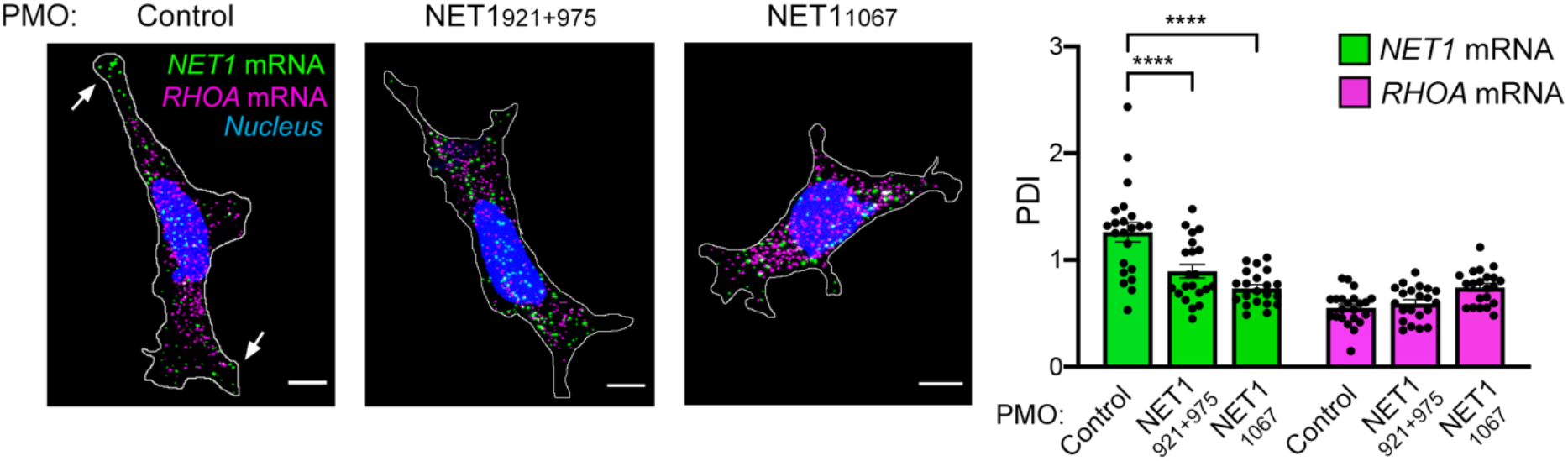
*NET1* mRNA localization in HEK293 cells depends on GA-rich sequences in the 3’UTR. Representative RNA FISH images of HEK293 cells treated for 72hrs with the indicated PMOs and PDI quantification. n=21 cells. Error bars: SEM. p-values: ****<0.0001 by one-way ANOVA. Arrows indicate *NET1* mRNA localized at peripheral protrusions. NET1-targeting PMOs prevent peripheral localization of *NET1* mRNA, but do not alter the distribution of *RHOA* mRNA. Scale bars: 10μm.

**Figure S6:**
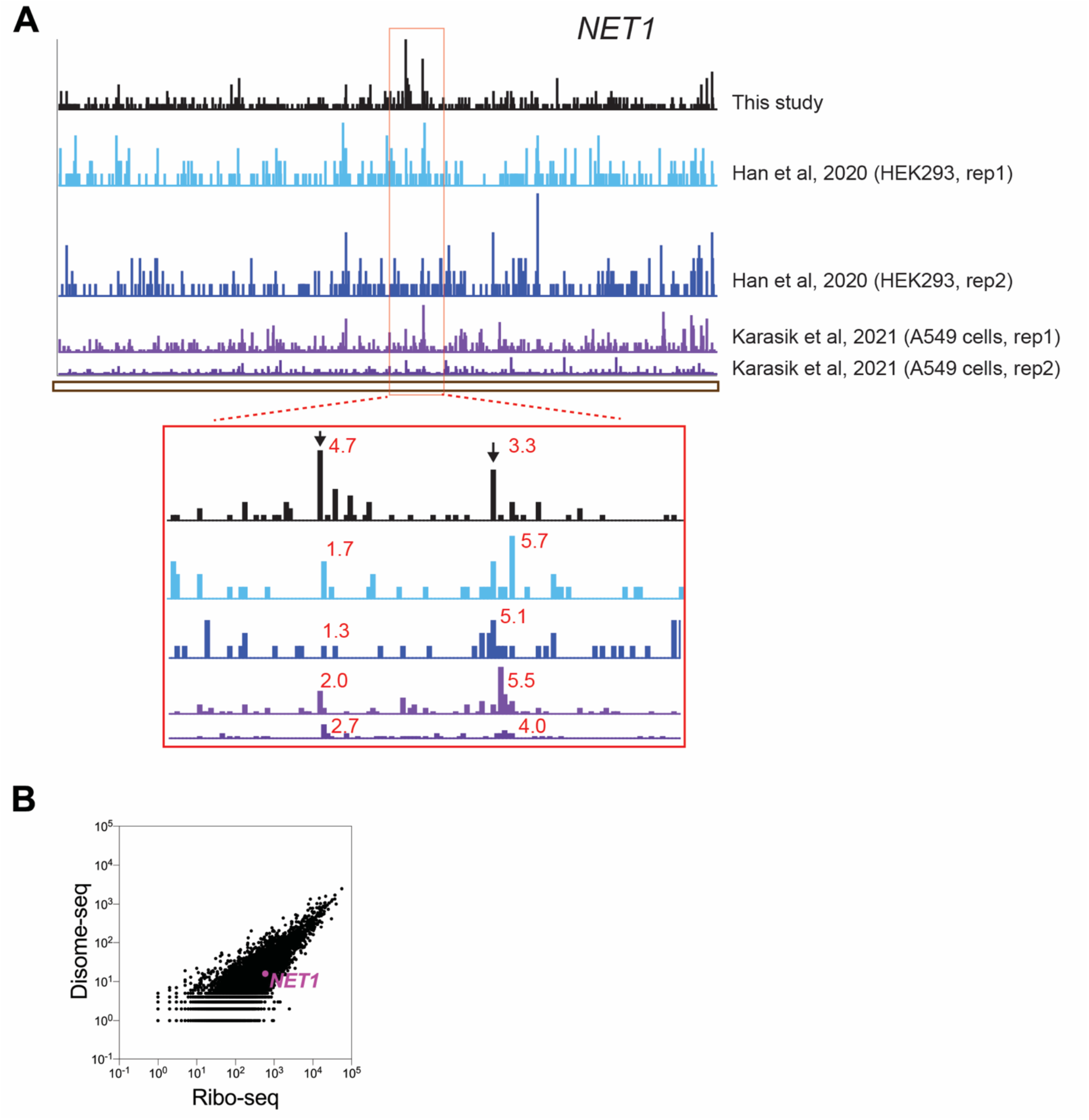
Ribosome profiling data of the NET1 transcript. **(A)** Snapshot of Ribo-seq reads mapped to the NET1 transcript in different datasets. Magnified inset at the bottom shows the reproducibility of both stalling peaks. The numbers in red next to each stalling peak indicate the pause score, which is calculated by averaging the reads -/+ 5 nt of the stalling peak and dividing that number to the average reads mapped to NET1 coding region. **(B)** Comparison of raw Disome-seq and Ribo-seq reads. NET1 is highlighted, where the number of Disome-seq and Ribo-seq reads mapped to NET1 are 16 and 591, respectively.

**Figure S7:**
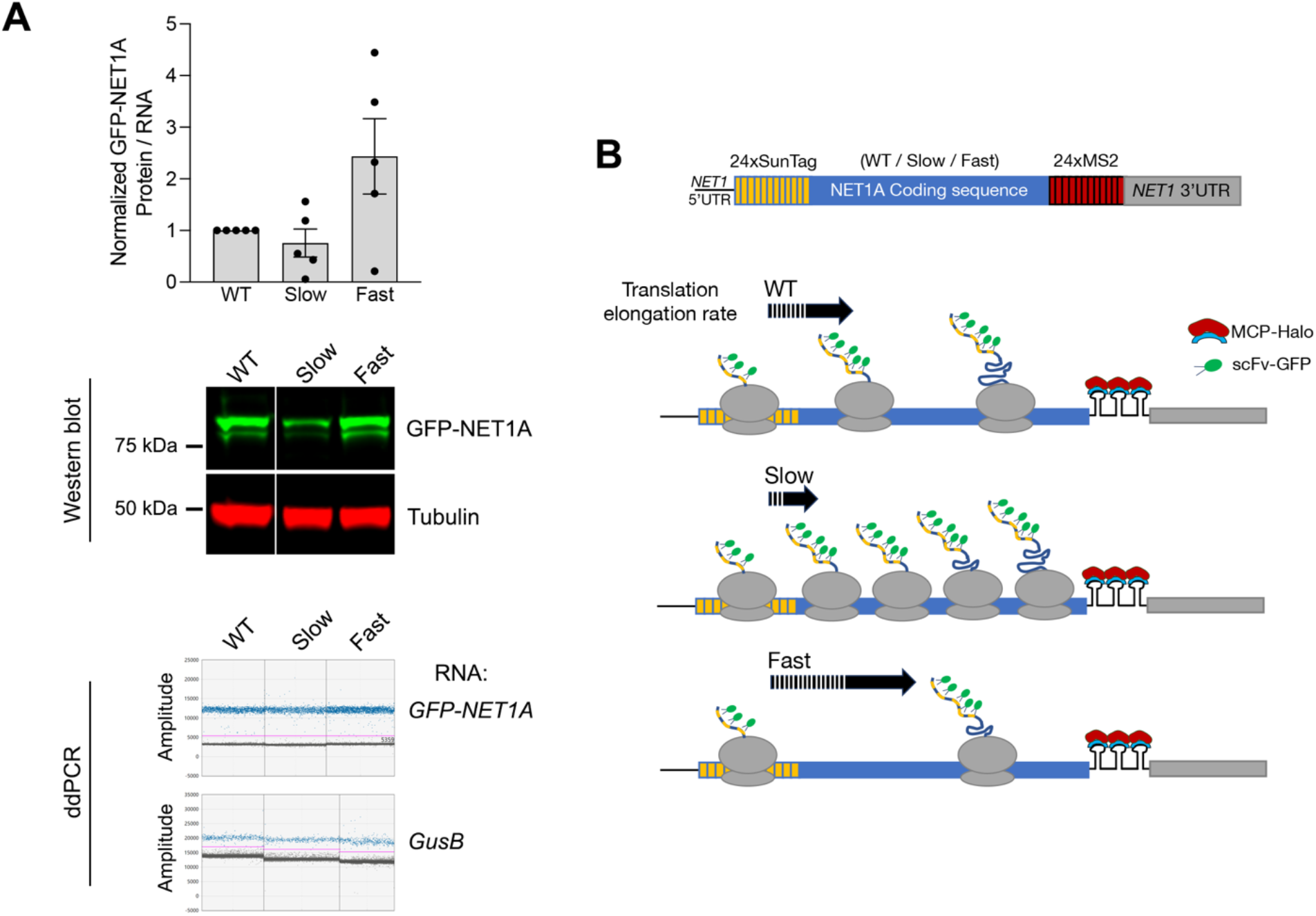
NET1A mutants with altered translation elongation rates. **(A)** Normalized GFP-NET1A protein/RNA ratio for the indicated stable GFP-NET1A-expressing cell lines (wild type (WT), slow or fast). Protein levels determined by Western blot; RNA levels by ddPCR. Representative results are shown in respective panels underneath the graph. Tubulin protein and *GusB* RNA were used as normalization controls. Error bars: SEM **(B)** Upper: schematic of overall structure of reporter constructs used for single-molecule translation site imaging. Reporters contain the 5’- and 3’-UTR sequences of human *NET1*, the wild type NET1 coding sequence (or slow and fast mutant variants), and 24 copies of SunTag epitopes and MS2 binding sites, used for fluorescence imaging detection. Lower: Schematic predictions of changes in ribosome occupancy per mRNA induced by the indicated changes in translation elongation rates.

**Figure S8:**
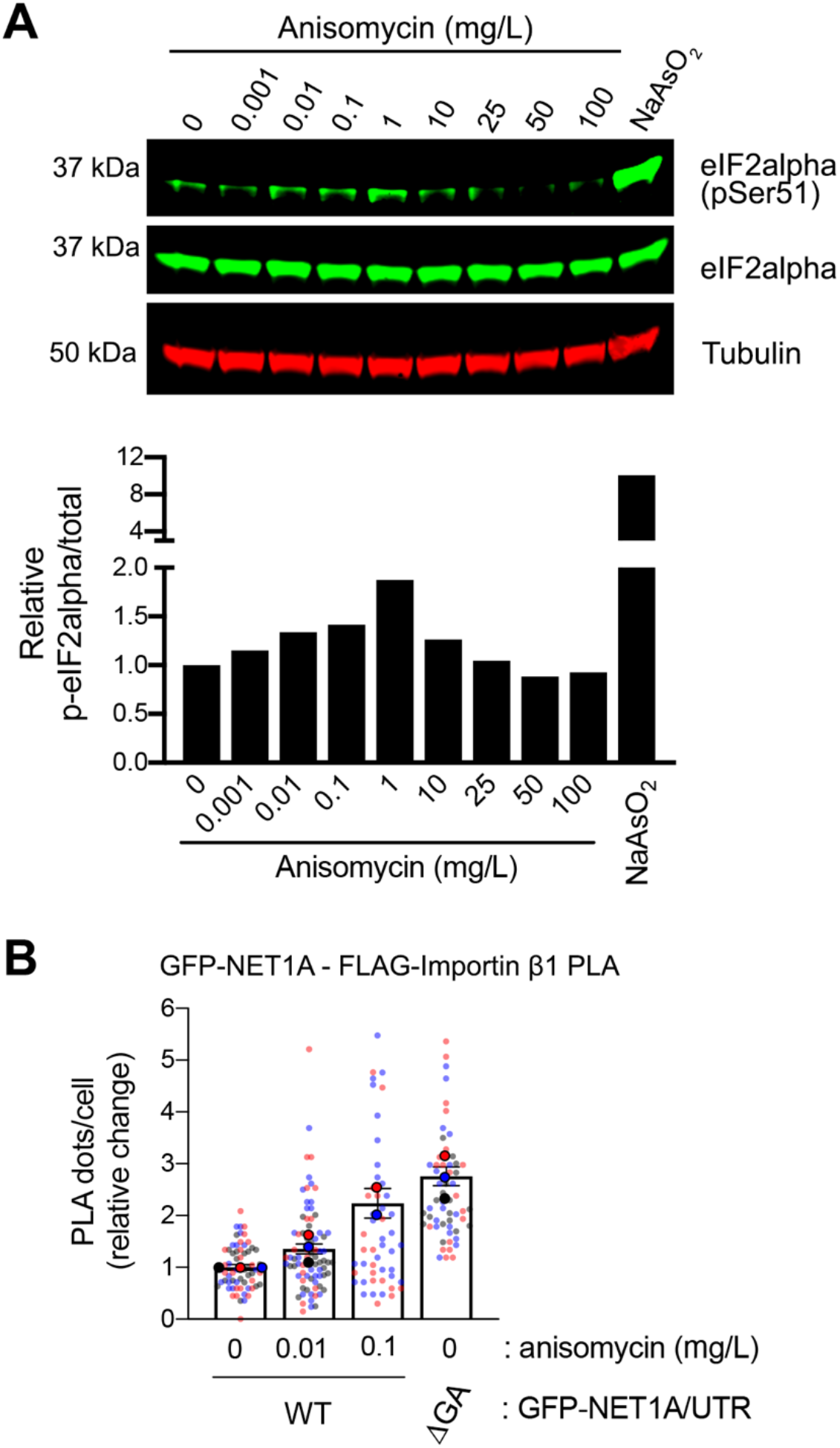
Sub-inhibitory concentrations of anisomycin increase interaction of NET1A with importin. **(A)** Western blot and quantification of pSer51-eIF2α levels in MDA-MB-231 cells treated with the indicated concentrations of anisomycin for 30min. For comparison, cells were treated with sodium arsenite for 30 min. **(B)** Quantification of in situ interaction between GFP-NET1A and FLAG-importin β1, by PLA of the indicated cell lines expressing GFP-NET1A from a peripheral (WT) or perinuclear RNA (ΔGA). n=52-80 in 2-3 independent experiments. In superplot, data points from individual replicates are color coded, and large, outlined color dots indicate the mean of each replicate. Error bars: SEM.

**Figure S9:**
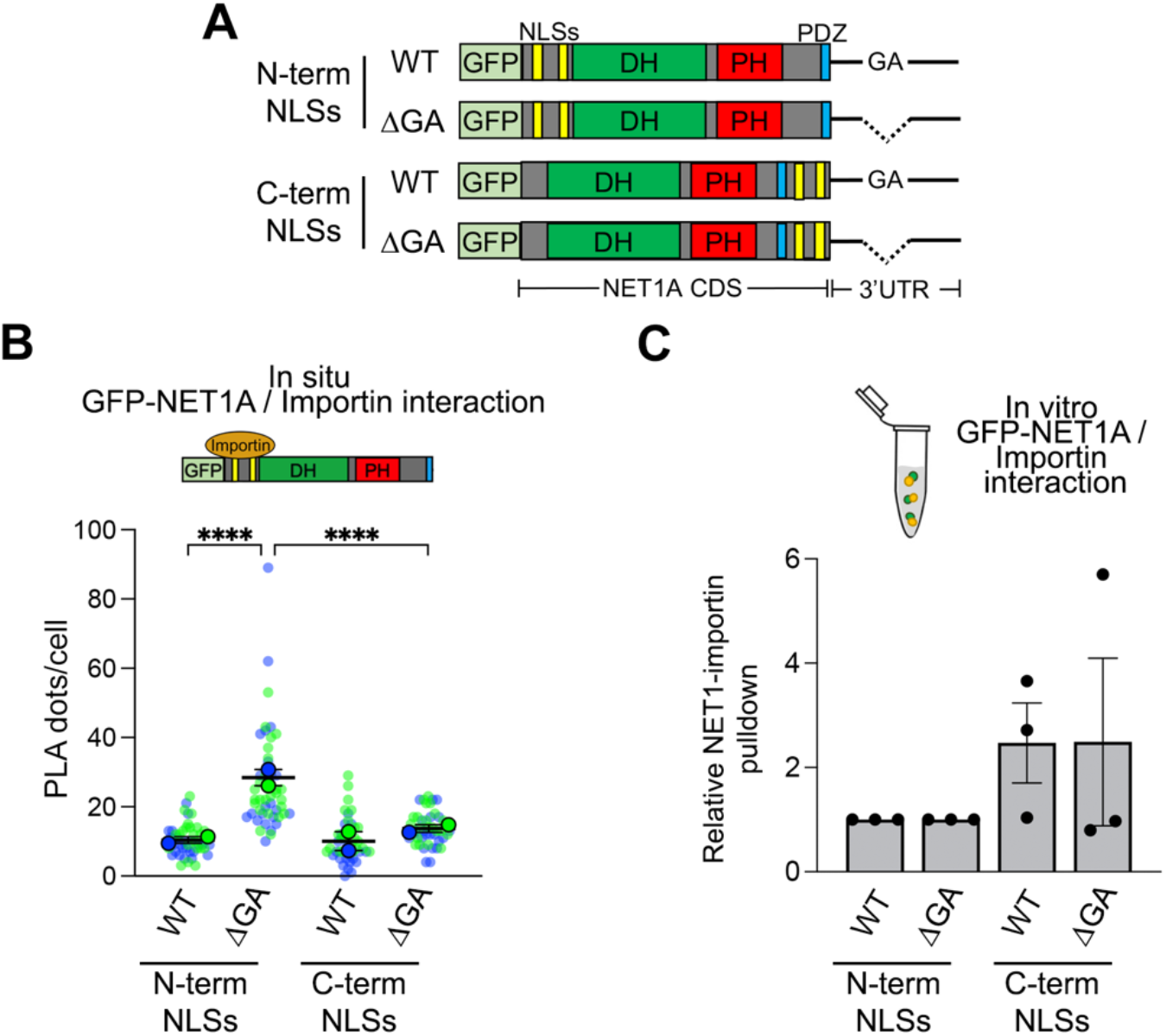
Order of NLS appearance influences NET1-importin interaction. **(A)** Schematic of GFP-NET1A constructs used for generation of stably expressing MDA-MB-231 cell lines. The normally occurring N-terminal NLSs were transferred to the C-terminus, thus being synthesized after the PH domain. Variants were expressed from transcripts carrying either the WT 3’UTR or a mutant with a deletion of the GA-rich region. **(B)** Quantification of in situ interaction between GFP-NET1A and FLAG-importin β1, by PLA of the indicated cell lines. n=45-49 from 2 independent experiments. **(C)** Quantification of relative pulldown efficiency of GFP-NET1A from lysates of the indicated cell lines with GST-importin α5. n=2. In superplots, data points from individual replicates are color coded, and large, outlined color dots indicate the mean of each replicate. Error bars: SEM. p-values: ****<0.0001, one-way ANOVA.

**Figure S10:**
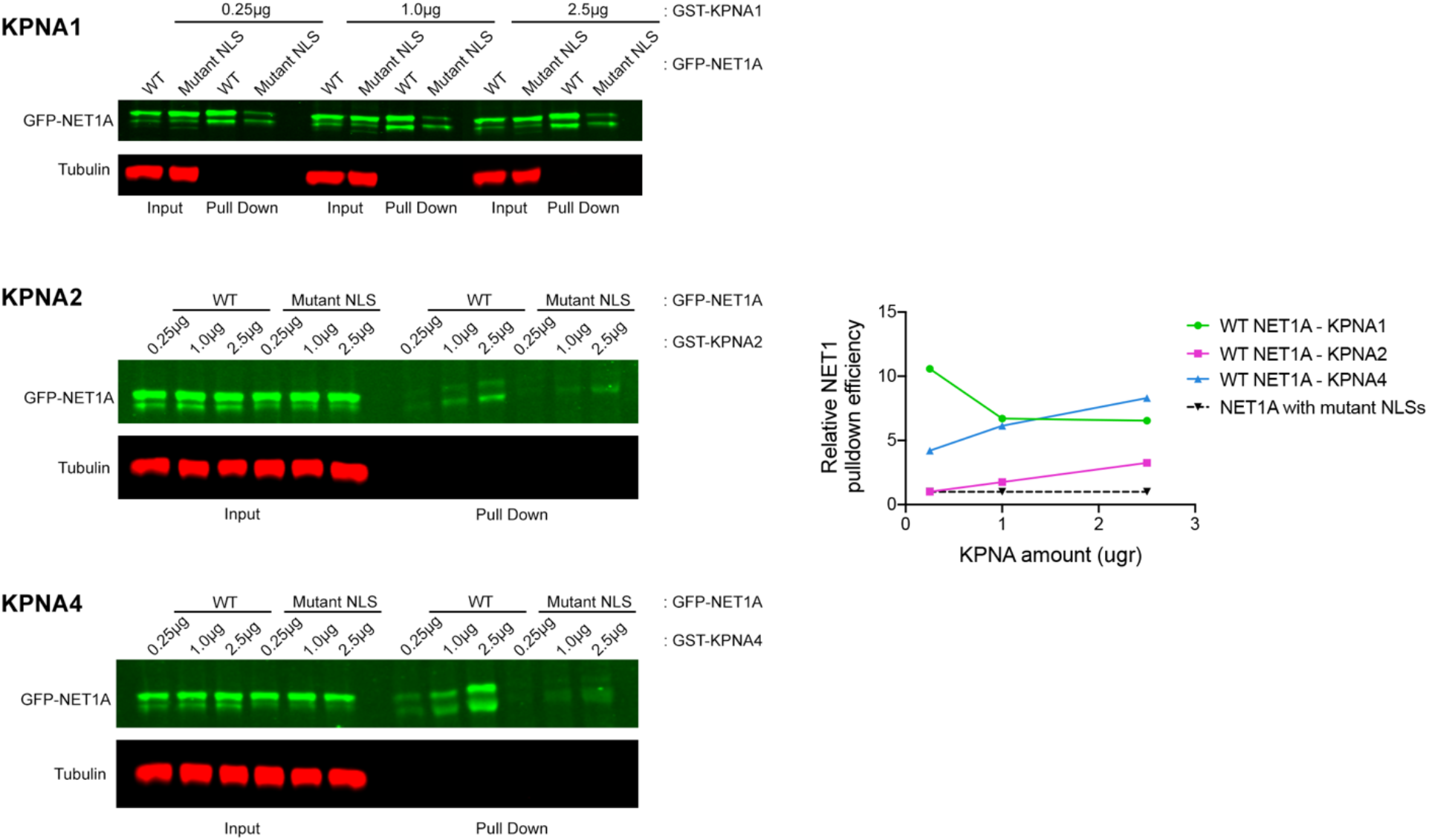
In vitro GFP-NET1A-importin α interaction. The indicated amounts of GST- importin α members (GST-KPNA1, GST-KPNA2, GST-KPNA4) were used for pulldown assays with lysates of GFP-NET1A-expressing cells. Pulldown efficiency is plotted relative to NLS mutant GFP-NET1A. GST-KPNA1 exhibits efficient binding at all concentrations tested and was used for further experiments.

**Figure S11:**
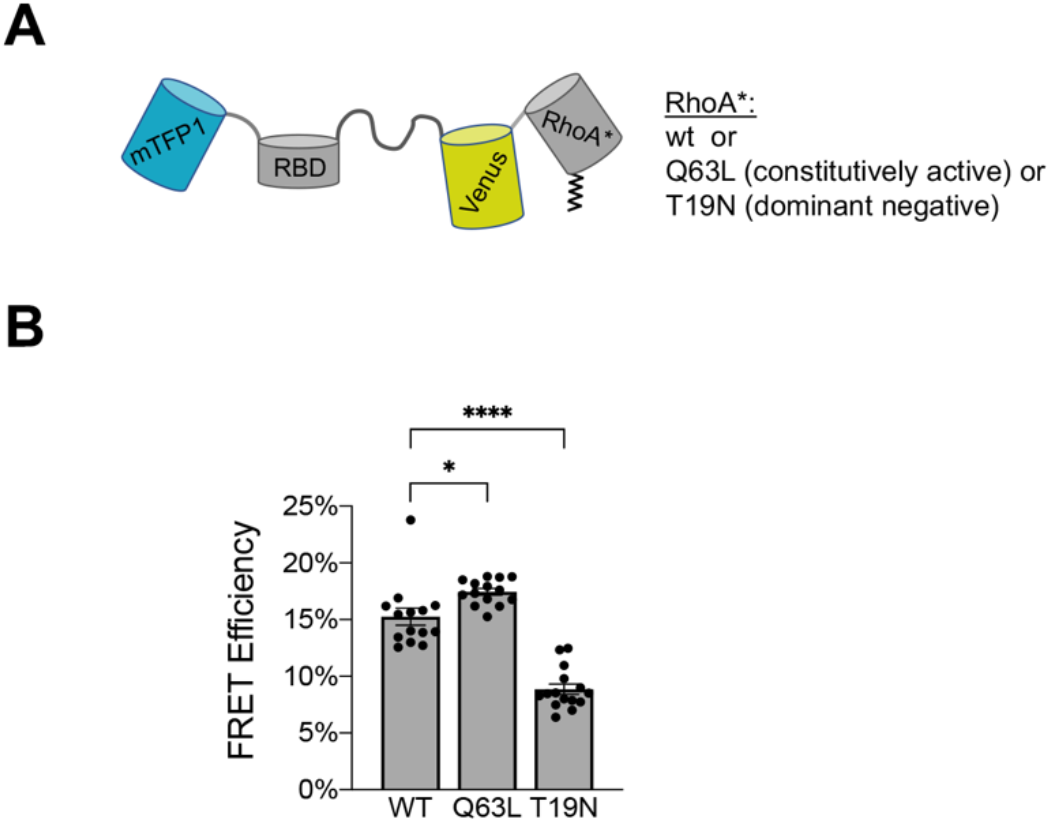
Validation of RhoA FRET biosensor. **(A)** Schematic of the biosensor used for RhoA activity estimation. **(B)** FRET efficiency measurements from a biosensor carrying wt RhoA, Q63L RhoA (constitutively active), or T19N RhoA (dominant negative). The biosensor can distinguish different levels of RhoA activity. Error bars: SEM. p-values: *<0.05, ****<0.0001 by one-way ANOVA.

**Figure S12:**
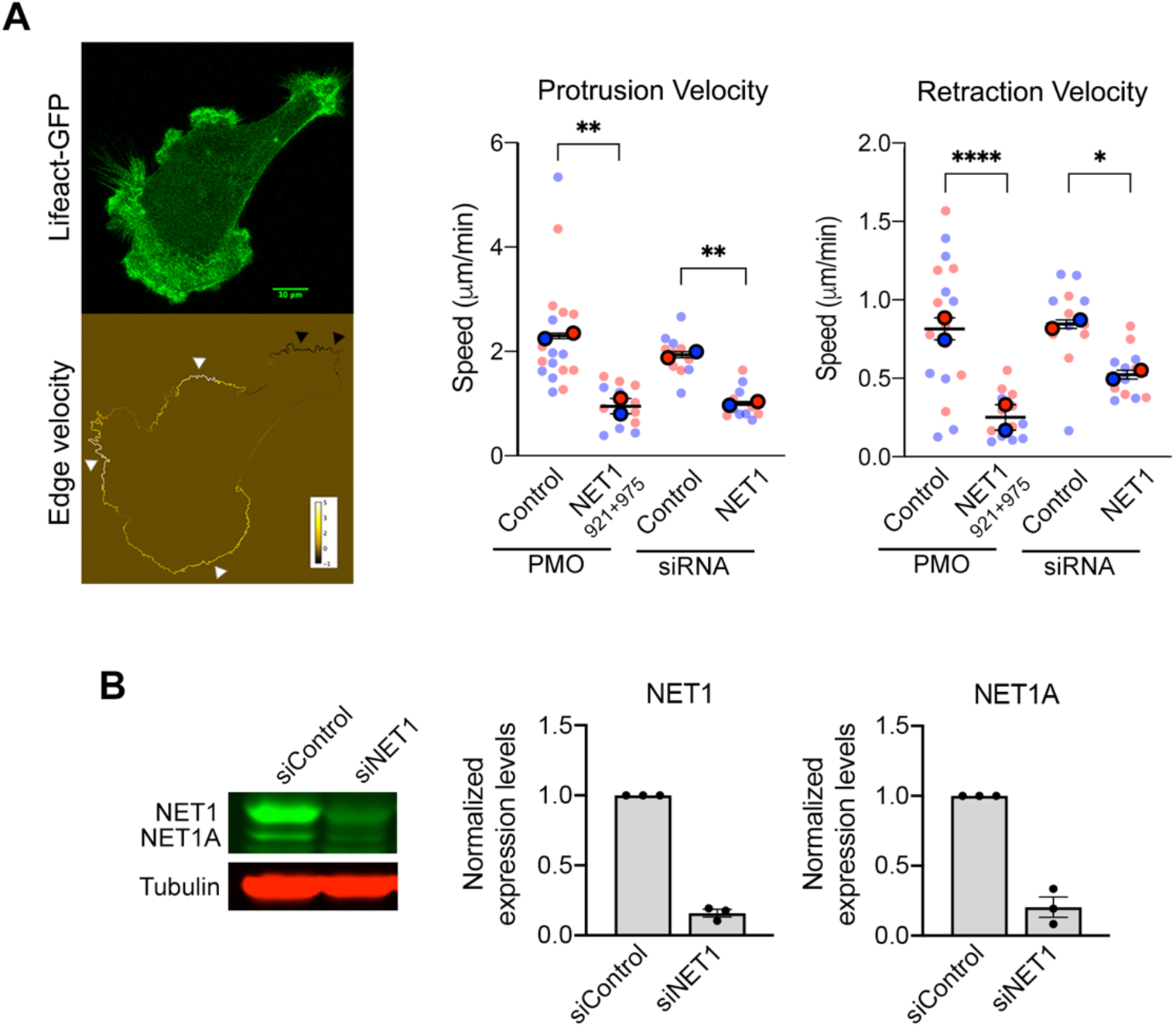
Peripheral *NET1* mRNA localization is essential for NET1 function in membrane edge dynamics. **(A)** Membrane protrusion and retraction speeds were measured in Lifeact-GFP cells treated with the indicated PMOs or siRNAs. Average protrusion and retraction velocities were calculated using an automated script. Image panels (left) show snapshots of time lapse movie (upper) and analysis output (bottom). The script measures edge velocity (protrusion: white arrowheads and retraction: black arrowheads) between consecutive frames of each time lapse movie and calculates average values for each analyzed cell. n=12-17 cells from two independent biological replicates. Error bars: SEM. p-values: *<0.05, **<0.01, ****<0.0001 by one-way ANOVA. Scale bar = 10um. **(B)** Western blot analysis of NET1 and NET1A protein levels upon siRNA treatment.

