## Supplementary material for "mRNA Location and Translation Rate Determine Protein Targeting to Dual Destinations": Table S1

**Table S1:** Sequences of coding region fragments used for generation of plasmid constructs

**WT NET1A:** Coding Sequence

gtggcacatgatgagactggaggtctcctacctattaaaaggaccatacgagtcctagatgtcaataaccagtccttcagagaacaaga
ggagccaagcaataaaagagttcgacctctggctcgtgtcacgtccttggcaaatttaatctctcctgtaagaaatggagctgtcagacgt
tttggtcaaacaatacagtcatttacccttcgtggtgaccacagatccccagcctctgcccagaagttttctagcaggtcaacagtcccaac
acccgccaagagaaggagcagtgcactgtggtcagagatgctggacatcaccatgaaggagtctctcaccaccagggagatcagacgg
caggaggcaatatatgaaatgtcccgaggtgaacaggatttaattgaggatctcaaacttgcaagaaaggcctaccatgaccccatgtt
aaagttgtccatcatgtcagaagaggaactcacacatatatttggtgatctggactcttacatacctctgcatgaagatttgttgacaaga
ataggagaagcaaccaagcctgatggaacagtggagcagattggtcacattctcgtgagctggttaccgcgcttgaatgcctacagagg
ttactgtagtaaccagctggcagccaaagctcttcttgatcaaaagaaacaggatccaagagtccaagacttcctccagcgatgtctcga
gtctcccttcagtcgaaaactagatctttggagtttcctagatatccctcgaagtcgcctagtcaaatacccttttactgttaaaagaaattct
taaacacactccaaaagagcaccctgatgttcagcttctggaggatgctatattgataatacagggagtcctctctgatatcaacttgaag
aaaggtgaatccgagtgccagtattacatcgacaagctggagtacctggatgaaaagcagagggaccccagaatcgaagcgagcaaa
gtgctgctgtgccatggggagctgcggagcaagagtggacataaactttacattttcctgtttcaagacatcttggttctgactcggcccgt
cacacggaacgaacggcactcttaccaggtttaccggcagccaatcccagtccaagagctagtcctagaagacctgcaggatggagatg
tgagaatgggaggctcctttcgaggagctttcagtaactcagagaaagctaaaaatatctttagaattcgcttccatgacccctctccagc
ccagtctcacactctgcaagccaatgacgtgttccacaagcagcagtggttcaactgtattcgagcggccattgccccccttccagtcggca
ggcagtccacctgagctgcagggcctgccggagctgcacgaagagtgtgaggggaaccacccctctgcgaggaaactcacagcccaga
ggagggcatccacagtttccagtgttactcaggtagaagttgatgaaaacgcttacagatgtggctctggcatgcagatggcagaggac
agcaagagcttaaagacacaccagacacagcccggcatccgaagagcgagggacaaagccctttctggtggcaaacggaaagagact
ttggtgtag

**Slow Mutant:** NET1A coding sequence shown (region highlighted contains mutations to introduce suboptimal codons)

gtggcacatgatgagactggaggtctcctacctattaaaaggaccatacgagtcctagatgtcaataaccagtccttcagagaacaaga
ggagccaagcaataaaagagttcgacctctggctcgtgtcacgtccttggcaaatttaatctctcctgtaagaaatggagctgtcagacgt
tttggtcaaacaatacagtcatttacccttcgtggtgaccacagatccccagcctctgcccagaagttttctagcaggtcaacagtcccaac
acccgccaagagaaggagcagtgcactgtggtcagagatgcttgacataacgatgaaggagtctttgacgacgagggagataagacgg
caggaggcgatatatgaaatgtcacgaggtgaacaagatttaattgaggatctaaaacttgcgagaaaggcgtaccatgacccgatgtt
aaagttgtccataatgtcagaagaggaactaacacatatatttggtgatctagactcttacatacctcttcatgaagatttgttgacaaga
ataggtgaagcgacgaagccggatggtacagtagagcaaataggtcacatactcgttagctggttaccgcgcttgaatgcgtacagagg
ttactgtagtaaccagttggcagccaaagctcttcttgatcaaaagaaacaggatccaagagtccaagacttcttgcagcgatgtttgga
gtctcccttcagtcgaaaactagatctttggagtttcctagatataccgcgaagtcgcctagtcaaatacccgttactgttaaaagaaattc
ttaaacacactccaaaagagcaccctgatgttcagcttctcgaggatgcgatattgataatacaaggagtcctctctgatataaacttgaa
gaaaggtgaatccgagtgccaatattacatagacaagcttgagtacctagatgaaaagcagagggaccccagaatagaagcgagcaa
agtgttgctttgccatggggagctgcggagcaagagtggacataaactttacattttcctgtttcaagacatcttggttctgactcggcccg
tcacacggaacgaacggcactcttaccaggtttaccggcagccaatcccagtccaagagctagtcctagaagacctgcaggatggagat
gtgagaatgggaggctcctttcgaggagctttcagtaactcagagaaagctaaaaatatctttagaattcgcttccatgacccctctccag
cccagtctcacactctgcaagccaatgacgtgttccacaagcagcagtggttcaactgtattcgagcggccattgccccccttccagtcggc

aggcagtccacctgagctgcagggcctgccggagctgcacgaagagtgtgaggggaaccacccctctgcgaggaaactcacagcccag
aggagggcatccacagtttccagtgttactcaggtagaagttgatgaaaacgcttacagatgtggctctggcatgcagatggcagagga
cagcaagagcttaaagacacaccagacacagcccggcatccgaagagcgagggacaaagccctttctggtggcaaacggaaagagac
tttggtgtag

**Fast mutant:** NET1A coding sequence shown (regions highlighted contain the mutations of the identified stall sites)

gtggcacatgatgagactggaggtctcctacctattaaaaggaccatacgagtcctagatgtcaataaccagtccttcagagaacaaga
ggagccaagcaataaaagagttcgacctctggctcgtgtcacgtccttggcaaatttaatctctcctgtaagaaatggagctgtcagacgt
tttggtcaaacaatacagtcatttacccttcgtggtgaccacagatccccagcctctgcccagaagttttctagcaggtcaacagtcccaac
acccgccaagagaaggagcagtgcactgtggtcagagatgctggacatcaccatgaaggagtctctcaccaccagggagatcagacgg
caggaggcaatatatgaaatgtcccgaggtgaacaggatttaattgaggatctcaaacttgcaagaaaggcctaccatgaccccatgtt
aaagttgtccatcatgtcagaagaggaactcacacatatatttggtgatctggactcttacatacctctgcatgaagatttgttgacaaga
ataggagaagcaaccaagcctgatggaacagtggagcagattggtcacattctcgtgagctggttaccgcgcttgaatgcctacagagg
ttactgtagtaaccagctggcagccaaagctcttcttgatcaaaagaaacaggatccaagagtccaagacttcctccagcgatgtctcga
gtctcccttcagtcgaaaactagatctttggagtttcctagatatccctcgaagtcgcctagtcaaaggcggcctgctgttaaaagaaattc
ttaaacacactccaaaagagcacggcggcgtgcagcttctggaggatgctatattgataatacagggagtcctctctgatatcaacttga
agaaaggtgaatccgagtgccagtattacatcgacaagctggagtacctggatgaaaagcagagggaccccagaatcgaagcgagca
aagtgctgctgtgccatggggagctgcggagcaagagtggacataaactttacattttcctgtttcaagacatcttggttctgactcggccc
gtcacacggaacgaacggcactcttaccaggtttaccggcagccaatcccagtccaagagctagtcctagaagacctgcaggatggaga
tgtgagaatgggaggctcctttcgaggagctttcagtaactcagagaaagctaaaaatatctttagaattcgcttccatgacccctctcca
gcccagtctcacactctgcaagccaatgacgtgttccacaagcagcagtggttcaactgtattcgagcggccattgccccttccagtcgg
caggcagtccacctgagctgcagggcctgccggagctgcacgaagagtgtgaggggaaccacccctctgcgaggaaactcacagccca
gaggagggcatccacagtttccagtgttactcaggtagaagttgatgaaaacgcttacagatgtggctctggcatgcagatggcagagg
acagcaagagcttaaagacacaccagacacagcccggcatccgaagagcgagggacaaagccctttctggtggcaaacggaaagag
actttggtgtag

**Mutant NLS:** NET1A coding sequence shown (regions highlighted contain mutations of amino acids within two basic NLSs):

gtggcacatgatgagactggaggtctcctacctattaaaaggaccatacgagtcctagatgtcaataaccagtccttcagagaacaaga
ggagccaagcaatgccgccgttgccctctggctcgtgtcacgtccttggcaaatttaatctctcctgtaagaaatggagctgtcagacgtt
ttggtcaaacaatacagtcatttacccttcgtggtgaccacagatccccagcctctgcccagaagttttctagcaggtcaacagtcccaac
acccgccgctgctgccagcagtgcactgtggtcagagatgctggacatcaccatgaaggagtctctcaccaccagggagatcagacggc
aggaggcaatatatgaaatgtcccgaggtgaacaggatttaattgaggatctcaaacttgcaagaaaggcctaccatgaccccatgtta
aagttgtccatcatgtcagaagaggaactcacacatatatttggtgatctggactcttacatacctctgcatgaagatttgttgacaagaat
aggagaagcaaccaagcctgatggaacagtggagcagattggtcacattctcgtgagctggttaccgcgcttgaatgcctacagaggtt
actgtagtaaccagctggcagccaaagctcttcttgatcaaaagaaacaggatccaagagtccaagacttcctccagcgatgtctcgagt
ctcccttcagtcgaaaactagatctttggagtttcctagatatccctcgaagtcgcctagtcaaataccctttactgttaaaagaaattctta
aacacactccaaaagagcaccctgatgttcagcttctggaggatgctatattgataatacagggagtcctctctgatatcaacttgaaga
aaggtgaatccgagtgccagtattacatcgacaagctggagtacctggatgaaaagcagagggaccccagaatcgaagcgagcaaag
tgctgctgtgccatggggagctgcggagcaagagtggacataaactttacattttcctgtttcaagacatcttggttctgactcggcccgtc
acacggaacgaacggcactcttaccaggtttaccggcagccaatcccagtccaagagctagtcctagaagacctgcaggatggagatgt

gagaatgggaggctcctttcgaggagctttcagtaactcagagaaagctaaaaatatctttagaattcgcttccatgacccctctccagcc
cagtctcacactctgcaagccaatgacgtgttccacaagcagcagtggttcaactgtattcgagcggccattgccccttccagtcggcag
gcagtccacctgagctgcagggcctgccggagctgcacgaagagtgtgaggggaaccacccctctgcgaggaaactcacagcccagag
gagggcatccacagtttccagtgttactcaggtagaagttgatgaaaacgcttacagatgtggctctggcatgcagatggcagaggaca
gcaagagcttaaagacacaccagacacagcccggcatccgaagagcgagggacaaagccctttctggtggcaaacggaaagagactt
tggtgtag

**C-term NLS:** NET1A coding sequence shown (region highlighted contains NLSs (green) and was transferred from the N-terminus to the C-terminus of the protein):

tggtcagagatgctggacatcaccatgaaggagtctctcaccaccagggagatcagacggcaggaggcaatatatgaaatgtcccgag
gtgaacaggatttaattgaggatctcaaacttgcaagaaaggcctatcatgaccccatgttaaagttgtccatcatgtcagaagaggaac
tcacacatatatttggtgatctggactcttacatacctctgcatgaagatttgttgacaagaataggagaagcaaccaagcctgatggaac
agtggagcagattggtcacattctcgtgagctggttaccgcgcttgaatgcctacagaggttactgtagtaaccagctggcagccaaagc
tcttcttgatcaaaagaaacaggatccaagagtccaagacttcctccagcgatgtctcgagtctcccttcagtcgaaaactagatctttgg
agtttcctagatatccctcgaagtcgcctagtcaaatacccttactgttaaaagaaattcttaaacacactccaaaagagcaccctgatg
ttcagcttctggaggatgctatattgataatacagggagtcctctctgatatcaacttgaagaaaggtgaatccgagtgccagtattacat
cgacaagctggagtacctggatgaaaagcagagggaccccagaatcgaagcgagcaaagtgctgctgtgccatggggagctgcggag
caagagtggacataaactttacattttcctgtttcaagacatcttggttctgactcggcccgtcacacggaacgaacggcactcttaccag
gtttaccggcagccaatcccagtccaagagctagtcctagaagacctgcaggatggagatgtgagaatgggaggctcctttcgaggagc
tttcagtaactcagagaaagctaaaaatatctttagaattcgcttccatgacccctctccagcccagtctcacactctgcaagccaatgac
gtgttccacaagcagcagtggttcaactgtattcgagcggccattgccccttccagtcggcaggcagtccacctgagctgcagggcctgc
cggagctgcacgaagagtgtgaggggaaccacccctctgcgaggaaactcacagcccagaggagggcatccacagtttccagtgttact
caggtagaagttgatgaaaacgcttacagatgtggctctggcatgcagatggcagaggacagcaagagcttaaagacacaccagacac
agcccggcatccgaagagcgagggacaaagccctttctggtggcaaacggaaagagactttggtgGTCGACggaccagtcgcactc
GTGGCACATGATGAGACTGGAGGTCTCCTACCTATTAAAAGGACCATACGAGTCCTAGATGTCAATAAC
CAGTCCTTCAGAGAACAAGAGGAGCCAAGCAATAAAAGAGTTCGACCTCTGGCTCGTGTCACGTCCTTG
GCAAATTTAATCTCTCCTGTAAGAAATGGAGCTGTCAGACGTTTTGGTCAAACAATACAGTCATTTACCC
TTCGTGGTGACCACAGATCCCCAGCCTCTGCCCAGAAGTTTTCTAGCAGGTCAACAGTCCCAACACCCGC
CAAGAGAAGGAGCAGTGCACTGTAA
