## Supplementary material for "mRNA Location and Translation Rate Determine Protein Targeting to Dual Destinations": Table S2

**Table S2:** Sequences of morpholino oligonucleotides (PMOs)

| PMO name | Sequence |
|---|---|
| Control | cctcttacctcagttacaatttata |
| NET1-16 | aacaaacagtctctcccatcagtta |
| NET1-68 | gtaaccctatgatgctccccttacg |
| NET1-146 | atctgtatatctttagctgccatta |
| NET1-219 | ataccaggcttaccatgttctaaat |
| NET1-354 | actggcagaattccagcatttgcaa |
| NET1-579 | aaataatcggcaggttaaatgaatg |
| NET1-608 | gctaaaaactactttacaataaaaa |
| NET1-656 | agacatgccaatttgaaaaggcatc |
| NET1-681 | ttaatcaggaagaaaaatactttaa |
| NET1-711 | acacacacacacacacatacacaca |
| NET1-761 | cttggtttcacttggtaaaattaat |
| NET1-796 | tggcaattttcttaattggctcaaa |
| NET1-828 | ggtctttaccctgaaatgctacact |
| NET1-853 | ctagaatacatcaagccatttcatg |
| NET1-878 | tttgaagtggttttctttcagtagt |
| NET1-921 | tccctcttgcatttcagacaacact |
| NET1-975 | gacaaaactactctcttttcctctc |
| NET1-1003 | tctggcacaaccagacattttactt |
| NET1-1042 | agttgagcttctcctatctcctttc |
| NET1-1067 | ttctacaacttactacacgccctca |
| NET1-1097 | aaggcaaataagtccacgtcccctc |
| NET1-1130 | tttcaaactcattatttgcaggtat |
| NET1-1169 | tctaatatggtcaaatttttaacac |
| NET1-1235 | gaaacattttgtaataaaagattca |
| NET1-1273 | tcctccccccttttcaaagatgatga |
| NET1-1324 | atagtcaacactgacttgaattgat |
| NET1-1351 | ttccactggccaaatatatttcaca |
| NET1-1386 | ggatggatctatttacagtcttttc |
| NET1-1411 | attcatttgtacagagaaatcattt |
| NET1-1449 | gtgattatgtgtgtgcttttttttt |
